# Interplay of diffusive and active motion governs cohesin extrusion and CTCF barrier function

**DOI:** 10.64898/2026.11.10.765814

**Authors:** Maxime M.C. Tortora, Geoffrey Fudenberg

## Abstract

Cohesin-mediated loop extrusion constitutes a crucial process for the regulation of chromosome folding and gene expression. A rapidly-growing body of *in vitro* experiments suggests that cohesin molecular motor activity is finely modulated by both chemical and mechanical cues, whose interplay holds important implications in living cells for development and disease. Yet, biophysical models capable of accurately and efficiently capturing the breadth of reported cohesin behaviors, along with their impacts on DNA spatio-temporal organisation, remain lacking. Here, building on multiple experimental observations of single-cohesin dynamics, we introduce a mesoscale description of cohesin based on the general framework of jump-diffusion processes. We show that the finite stall force of cohesin naturally emerges from the conjunction of active, directional extrusion moves with passive, stochastic drift. The tension-dependent competition between these two orthogonal modes of motion gives rise to highly-heterogeneous cohesin kinetics featuring extrusion, translocation, and loop slipping phases, as observed *in vitro*. These behaviors are predicted to depend sensitively on both buffer composition and nucleosomal content, reconciling the surprising variability of cohesin kinetics reported across experimental conditions. This jump-diffusion model (JDM) is fully parametrized using available single-molecule measurements and bridges the timescale gap between microscopic stepping events and cohesin regulator association, thus providing mechanistic insights into how CTCF impedes extrusion. Hence, the JDM quantitatively describes force-dependent extrusion in both DNA curtain assays *in vitro* and in the context of native chromatin for genome folding predictions *in nucleo*, bridging these previously disparate contexts under a unified numerical and theoretical formalism.

**Significance statement:** Cohesin folds chromosomes into loops that help regulate gene expression, yet its diverse behaviors and sensitivity to mechanical forces remain largely unexplained. Here, we develop a theory for how competition between active DNA capture and passive sliding governs loop growth, shrinkage, and motion. This framework reconciles seemingly conflicting experimental observations, and predicts how weak forces generated by cohesin on naked DNA *in vitro* translate into tensions strong enough to drive large-scale genome folding on native chromatin. Jointly considering cohesin mechanics and biochemistry further enables us to explain how the DNA-binding protein CTCF modulates extrusion activity. By connecting single-molecule measurements with chromosome organization mechanisms, we thus provide a quantitative foundation for how physical conditions and regulatory proteins collectively shape genome folding.

## Introduction

Loop extrusion by Structural Maintenance of Chromosomes (SMC) complexes is a deeply conserved molecular mechanism for DNA organization (1, 2). The SMC family includes cohesin, condensin and SMC5/6, which share a common molecular structure: two SMC subunits with long coiled-coils connecting a hinge and globular ATPase head domains, and a kleisin that bridges the two head domains. Extrusion by SMCs can occur either on naked DNA or chromatin, requires ATP hydrolysis, and can be regulated by additional transiently-associating protein cofactors. In mammalian interphase, the primary extruder cohesin requires association of the regulator NIPBL to be competent for loop formation (3, 4), and is blocked at sequence-specific positions by CTCF, a DNA-binding 11-zinc finger protein (5, 6). The ability of cohesin to extrude loops was first inferred indirectly by comparing polymer models with genomic data (7–9), and has since been corroborated by single-molecule experiments (4, 10–17). Beyond confirming the crucial molecular components for cohesin extrusion, *in vitro* assays revealed a surprisingly-rich range of kinetic behaviors that are highly-sensitive to detailed reconstitution conditions. Still, the detailed mechanisms governing heterogeneous cohesin behaviors and their implications in genomic contexts remain unclear.

To reconstitute loop extrusion *in vitro*, fixed-length DNA is typically labeled with intercalating dye and tethered by one or both ends to a glass slide (18). Cohesin and its cofactors, potentially conjugated with fluorescent labels, are then introduced through buffer flow controlled via microfluidics. This flow is usually maintained in single-tethered setups to physically stretch DNA for visualization, but may be switched off prior to data collection in double-tethered assays. Tether geometry and buffer flow thus jointly set the mechanical conditions of the assay, and images of labeled DNA, alongside tagged cohesin or cofactors, are then acquired over timescales of a few minutes. Trajectories from cohesin reconstitution assays reported rapid formation of DNA loops, with varying degrees of asymmetry. Closer analysis of these trajectories uncovered multiple kinetic behaviors, including loop growth, shrinkage, translocation, and diffusion (11, 16). During translocation and diffusion, loop size remains approximately constant while the loop moves directionally or stochastically along DNA. Loop growth is closely associated with NIPBL-cohesin colocalization, and NIPBL exchange can accompany changes in extrusion direction (16).

Mechanical conditions also influence these trajectories: DNA tension can stall net loop growth at low forces (10–12), and alter the apparent symmetry of extrusion (15). A theory for how tension and cofactor association collectively drive cohesin kinetics is currently at large.

Beyond displaying heterogeneous dynamics, cohesin behavior is sharply affected by changes in buffer and substrate composition. Both high-salt washes and ATP depletion can destabilize extruded loops (10, 11). Cohesin diffusion appears remarkably sensitive to ionic content (see Dataset S1) as well as the inclusion of nucleosomes and other nano-to-microscale obstacles onto DNA (19). This high susceptibility suggests diffusivity as an important potential regulator of SMC kinetics. Yet, how passive motion combines with active, ATP-dependent DNA capture to determine loop growth versus loss under varying tension, buffer and substrate conditions remains unresolved.

CTCF provides an exemplary case for how molecular interactions and mechanical conditions jointly dictate cohesin behavior. CTCF can block cohesin-mediated loop extrusion (6), with proposed mechanisms ranging from direct cohesin binding, to ATPase inhibition, to altered cofactor association (5). Yet CTCF barrier activity also depends strongly on DNA tension: in reconstituted assays, blocking is weak at low tensions and only becomes effective at higher tensions approaching the cohesin stall force (14).

Explaining why tension-dependent blocking emerges for CTCF remains a key theoretical challenge.

A unified understanding of cohesin extrusion thus requires modelling its joint susceptibility to chemical, physical and mechanical cues (20) over timescales of multiple minutes. However, most mesoscale models to date have either ignored local mechanics (21) or chemistry (15, 22–25). Conversely, other studies have focused on the molecular details of SMC stepping (26–29). Still, despite recent progress (30, 31), detailed models remain challenging to parametrize unambiguously using available experimental data, and are precluded by high computational costs from the multi-minute timescales over which cohesin binds CTCF, associates with its cofactors, and ultimately drives chromosome folding (21). Thus, new mesoscale approaches are needed to bridge the gap between single-molecule experiments *in vitro* and their *in vivo* implications.

Here we derive a jump-diffusion model (JDM) of cohesin to determine how the combination of active DNA capture and diffusive motion governs cohesin extrusion kinetics under tension. This framework reveals why cohesin is highly sensitive to changes in buffer and substrate conditions, and may be fully parametrized using available *in vitro* data. The model identifies force-biased diffusion as a source of loop slipping, shows its interplay with capture generates the full breadth of heterogeneous trajectory kinetics reported *in vitro*, and predicts that higher stall forces could emerge in native chromatin. The JDM further reveals that CTCF stalls cohesin through the combination of polar and non-polar pathways, thus providing a dual mechanistic basis for CTCF barrier functionality. Our general formalism may be readily extended to other SMCs beyond cohesin, and illustrates how single-molecule measurements of SMC kinetics may be leveraged into quantitative biophysical and biochemical models suitable for large-scale genome folding simulations.

## Results

### I. Tension-dependent cohesin motion emerges from jump-diffusion dynamics

Two key features of extrusion shared by many SMCs motivate our mesoscale model of cohesin. First, single-molecule micromanipulation by magnetic tweezers evidences that ATP-dependent DNA capture steps are only weakly affected by applied DNA tension and have lengths ϵ ≃ 40 *nm* (∼130 *bp*) (14, 32), slightly shorter than the dimensions of stretched SMC coils (33–36). This indicates that applied forces should primarily modify stepping rates, rather than step sizes. Second, trajectory analysis reveals that cohesin (10, 37), condensin (38) and SMC5/6 (39) also perform rapid, ATP-independent diffusion *in vitro* at high-enough salt concentrations. This highly-conserved diffusive behavior can be measured directly from single-molecule trajectories in various experimental conditions (19, 40, 41). Our theory connects these two modes of motion to three experimentally-accessible mesoscale quantities, namely: stall force (*F*_*stall*_), average extrusion velocity (<u>*v*</u>), and diffusivity (*D*). As a baseline, we considered: (i) median stall force *F*_*stall*_ ≃ 0.15 *pN*, as reported for both purified human cohesin (14) and Xenopus cohesin in whole cell extract (12); (ii) mean extrusion velocity <u>*v*</u> = 1 *kb*/*s* measured in purified HeLa (11) and fission yeast cohesin assays (13) and (iii) diffusivity 2*D* ≃ 0.01 *µm*^2^/*s*, estimated in standard extrusion buffer conditions (pH 7, ∼50 mM NaCl (10, 16); c.f. Dataset S1). Together, these enabled us to exactly parameterize a model that describes the sensitivity of SMC kinetics to substrate and buffer composition at the mesoscale.

Given the two modes of SMC motion, we modeled cohesin kinetics as driven both by active extrusion steps (“jumps”) and passive local diffusive motion. In this jump-diffusion model (JDM), individual cohesins were described as a set of two spatially-proximal “anchors”, whose positions define the base of a DNA loop (Fig. 1a). We considered NIPBL-bound cohesin to possess an active anchor on one side — subject to both ATP-independent diffusion and ATP-dependent DNA capture — and a passive anchor on the other, subject to diffusion alone (Fig. 1a). Given the reported insensitivity of cohesin diffusivity on NIPBL binding (16), we assumed diffusion kinetics were equivalent at passive and active anchors. To analytically explore the JDM, we first assumed all loaded cohesins to be NIPBL-bound with one active anchor, and later relaxed this assumption and explicitly modeled NIPBL exchange in Secs. III and IV. Applied DNA tension *F* separately affects both active and passive stepping rates, since reeling a DNA segment into a loop requires mechanical work commensurate with the segment length — while transferring DNA out of the loop results in a net energy gain (c.f. Sup. Sec. I). We thus considered that each cohesin anchor displaced via forward or backward moves (Fig. 1a), respectively resulting in loop growth or shrinkage, and derived how force separately modulates the four associated stepping rates: forward diffusion (*d*_*f*_), backward diffusion (*d*_*b*_), forward extrusion (*e*_*f*_) and backward extrusion (*e*_*b*_).

**Figure 1.**
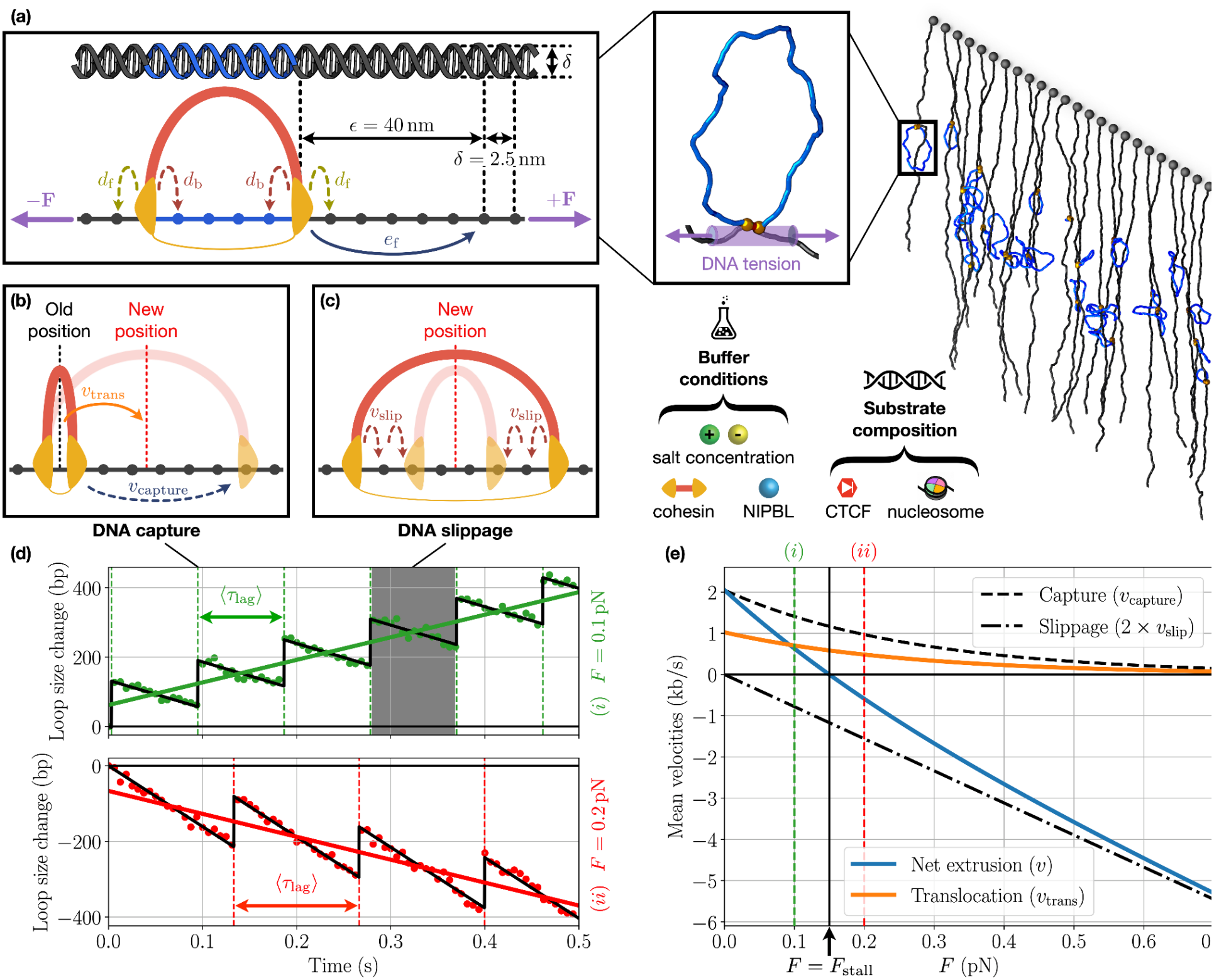
Tension-dependent cohesin motion emerges from jump-diffusion dynamics. **(a)** *Right*: Illustration of loop extrusion visualized in a typical in vitro reconstitution assay, where cohesin and cofactors are introduced to tethered DNA. Extruded loops are shown in blue, and the relevant portion of the DNA for tension shown in purple (inset). *Left:* The JDM features local (backward and forward) passive diffusion moves at both anchors with discrete forward extrusion steps of size ϵ = 40 *nm* at the active anchor. Their respective rates *d*_*b,f*_ and *e*_*f*_ are affected by applied tension *F*, substrate, and buffer conditions. **(b)-(c)** In the JDM, the extrusion velocity, quantifying loop growth versus loss, is governed by a competition between DNA capture by the active anchor (b) and DNA slippage at both the active and passive anchors (c). **(d)** Illustrative traces show how the staggered dynamics of capture (vertical dashed lines) and slippage (dots) set the net extrusion velocity (colored line slopes) yielding either loop growth (*i*, green) or loss (*ii*, red). The velocities of each process are determined by the tension-dependence of their respective rates *d*_*b,f*_ and *e*_*f*_ (Eqs. (2) and (4)). Note that both the time and the slippage velocity between DNA capture steps increase at higher forces. **(e)** Net capture (*v*_*capture*_, Eq. (4)), slippage (*v*_*slip*_, Eq. (2)), extrusion (*v*, Eq. (5)) and translocation (*v*_*trans*_, Eq. (8)) velocities resulting from the microscopic kinetics in Figs. 1a-d. Dashed vertical lines delineate the specific applied tension values illustrated in Fig. 1d.

For passive anchors, only the diffusive stepping rates are nonzero, and tension imposes a bias towards loop shrinkage. The functional form of this bias derives from the standard Boltzmann expression

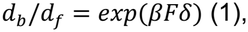

where β ≡ 1/*k*_*B*_*T* is the inverse temperature and δ the infinitesimal size of local diffusion steps. For small-enough δ, Eq. (1) leads to a force-dependent mean slippage velocity *v*_*slip*_ in the general fluctuation-dissipation form (see Supporting Information),

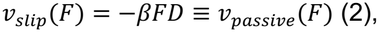

with *D* the one-dimensional diffusivity of loaded cohesins. Thus, in the absence of stretching forces, the ATP-independent motion of passive anchors is purely diffusive and stochastic [*v*_*slip*_(*F* = 0) = 0], while applied tension skews trajectories towards loop shrinkage [*v*_*slip*_(*F* > 0) < 0].

For active anchors, discrete ATP-dependent steps increase the loop size by ϵ in addition to passive slippage. The tension-induced relative bias in active stepping rates takes a similar form to Eq. (1),

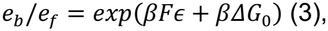

where Δ*G*_0_ < 0 corresponds to the free energy of ATP hydrolysis consumed in a single cohesin capture event, which promotes loop growth at low stretching forces (*e*_*b*_/*e*_*f*_ < 1 for *F* → 0). Eq. (3) thus enables estimation of the relative forward versus backward active stepping rates *e*_*f,b*_ from Δ*G*_0_, *F*, ϵ, and β (∼1/4.28 *pN*^−1^*nm*^−1^ at physiological temperature *T* = 310 *K*). For Δ*G*_0_, although cohesin features two distinct ATPase heads and could consume two ATP molecules per active step, we conservatively use the typical energy of a single ATP binding/hydrolysis event ( Δ*G*_0_ ≃ −50 *pN nm* (42)). Using *F*_*stall*_ ≃ 0.15 *pN*, along with the cohesin step size ϵ ≃ 40 *nm* (∼130 *bp*), Eq. (3) yields *e*_*b*_/*e*_*f*_ < 10^−4^ in the loop extrusion regime (*F* < *F*_*stall*_). Therefore, the backward active stepping rate *e*_*b*_ is predicted to be negligible and DNA capture by cohesin largely acts as a unidirectional mechanism given the stall forces observed in reconstituted extrusion assays, although backward active steps could be expected to occur at higher force regimes (see Discussion). In the *continuum limit* of jump processes (43), the mean loop growth velocity *v*_*capture*,_ resulting from active stepping then takes the form (see Supp. Sec. I)

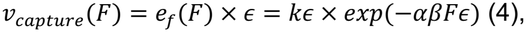

with *k* the cohesin stepping rate at zero tension and α ∈ [0,1] the Bell load distribution parameter, which describes the fraction of the applied force serving to inhibit forward power strokes (44). Thus, the net velocity *v*_*active*,_ ≡ *v*_*capture*,_ + *v*_*slip*_ of active anchors simply results from the interplay between active capture and passive slipping motion.

In the JDM, the mean loop extrusion velocity *v*(*F*) of a cohesin complex comes from the combined motion of both the cohesin active and passive anchors as a function of tension. Using Eqs. (2) and (4),

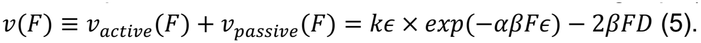

Using *v*(*F*_*stall*_) = 0, Eq. (5) provides a direct relation between cohesin stall force *F*_*stall*_, diffusion constant *D* and the *a priori* unknown model parameters *k* and α,

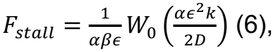

with *W*_0_ the principal branch of the Lambert *W* function. Eq. (5) additionally yields the mean observable cohesin extrusion velocity 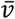, averaged over accessible tensions *F* < *F*_*stall*_,

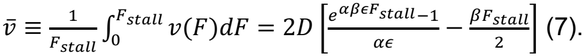

Note that Eq. (7) assumes that experimental tensions are uniformly distributed in the range *F* ∈ [0, *F*_*stall*_]; the quantitative accuracy of this approximation is explicitly validated in the Supporting Information (c.f. Fig. S1a). Hence, Eqs. (6) and (7) provide a set of two independent equations relating *k* and α to the experimentally-measurable quantities *F*_*stall*_, *D*, ϵ and 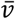 thus allowing the JDM to be fully constrained by available single-molecule data.

Using our baseline literature estimates for these parameters, Eqs. (6) and (7) yield α = 0.4 and *k* = 15.8 *s*^−1^. This computed α is consistent with those of other biomolecular motors (α ∼ 0.1–0.7 for myosin (45) and kinesin (46)), while the rate *k* implies that active steps should occur upwards of 15 times per second at low-enough applied tensions, comparable to orthogonal theoretical estimates (13). Such rapid DNA capture kinetics at low tensions are associated with cohesin loop extrusion velocities of up to *v* ≃ 2.2 *kb*/*s*, in remarkable agreement with the maximum values of 2.1–2.4 *kb*/*s* reported in DNA curtain assays (11, 13) (Fig. S1b).

The JDM also predicts that the active stepping rate *e*_*f*_(*F*) decays exponentially with increasing tensions (Eq. (4)), and may reach up to *e*_*f*_ = *k* ∼ 15 *s*^−1^ for *F* → 0, consistent with previous theoretical estimates (13, 30) but significantly higher than the rates of ∼1-2 ATP/cohesin/s reported by previous experimental studies (10, 11). This discrepancy can be explained by the fact that bulk ATPase assays cannot distinguish loaded, extrusive cohesins from unloaded or otherwise stalled populations, and thus generally only provide a lower bound for ATP hydrolysis kinetics (47). However, this active stepping rate drops to *e*_*f*_ ∼ 1 *s*^−1^ in the higher force regime *F* ∼ 0.5 *pN* typically accessible via magnetic tweezer micromanipulation (Fig. S1c), consistent in magnitude with experimental observations in such assays (14, 22).

More generally, the JDM predicts that the net extrusion velocity arises from the competition between active, one-sided DNA capture and passive DNA slippage (Fig. 1b,c; Movies S1-S3). This results in characteristic sawtooth-like kinetics of loop growth and shrinkage at the lengthscale of single DNA capture steps (ϵ ≃ 40 *nm* ∼ 130 *bp*; Fig. 1d). Importantly, tension increases both the lag time between DNA capture steps < τ_*lag*_ >= 1/*e*_*f*_(*F*) (Eq. (4)) and loop shrinkage velocity |*v*_*shrink*_| = 2β*FD* (Eq. (5)). For *F* < *F*_*stall*_, predicted loop shrinkage velocities (|*v*_*shrink*_| = 0–1.3 *kb*/*s*) closely align with experimental measurements (0–1.2 *kb*/*s* (16)). The JDM thus correctly predicts the presence of transient loop slipping windows even at low tensions conducive to extrusion. The brief duration of these slippage phases (τ_*lag*_ ∼ 100 *ms* in the active NIPBL-bound state) could be additionally modulated by cohesin biochemistry, as further discussed in Sec. III.

In the JDM, the alternation of DNA capture and slippage also results in the directional displacement of the complex — henceforth referred to as *translocation* — along the direction of cohesin DNA capture (Figs. 1b-c). The corresponding translocation velocity *v*_*trans*_ quantifying the motion of the loop position, defined as the geometric center of the loop (Figs. 1b-c), simply reads as

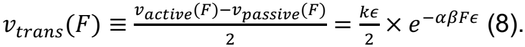

This results in typical velocities *v*_*trans*_(*F*) = 0.6–1 *kb*/*s* for *F* < *F*_*stall*_, in the force range where cohesin is competent for loop extrusion, but translocation interestingly still persists in the regime *F* > *F*_*stall*_, in which extrusion ability is fully inhibited (Fig. 1e). In these higher tension conditions, *v*_*trans*_ spans the predicted range *v*_*trans*_(*F*) = 0–0.6 *kb*/*s*, highly consistent with the values reported for cohesin translocation on strongly-stretched DNA (0–0.7 *kb*/*s* (11)). This value decays to zero only asymptotically at high forces, in which directional translocation gradually gives place to purely stochastic, undirected diffusion of the complex, as discussed in more detail in Sec. II.

Thus, the JDM recapitulates the full range of loop growth, shrinkage and translatory kinetics reported for cohesin in reconstituted assays, and provides a mechanistic, parameter-free framework that quantitatively captures tension-dependent extrusion on DNA *in vitro* at fixed buffer conditions.

### II. Cohesin behavior is determined by buffer and substrate conditions

Beyond their DNA tension-dependence, cohesin kinetics also appear highly sensitive to buffer conditions in reconstituted assays (10, 11, 16). Specifically, a broad range of cohesin diffusivities have been reported *in vitro* upon changes in DNA substrate composition (19), salt (19, 41) or cohesin regulator content (40). We thus tested whether the JDM generates the same dependence of extrusion kinetics on diffusivity as seen experimentally.

The JDM predicts the cohesin diffusivity *D* affects extrusion behavior by modulating the loop-slipping velocity (*v*_*slip*_, Eq. (2)), but leaves the cohesin capture velocity (*v*_*capture*,_ Eq. (4)) and the ensuing translocation velocity invariant (*v*_*trans*_, Eq. (8)). We thus varied *D* systematically in the range 2*D* ∼ 10^−4^– 1 *µm*^2^/*s* centered on our baseline (2*D* ∼ 10^−2^ *µm*^2^/*s, F*_*stall*_ = 0.15 *pN*; c.f. Sec. I), which encapsulates most commonly-reported experimental estimates (Dataset S1), and computed the corresponding extrusion velocities (Eq. (5)). This analysis revealed that the stall force *F*_*stall*_, which marks the threshold between loop growth and loss, depends strongly on diffusivity (Fig. 2a). Rapid diffusion suppressed evidence of extrusion activity by promoting loop slipping, with stall forces dropping to nearly zero at sufficiently high diffusivity 2*D* ∼ 1 *µm*^2^/*s* (Fig. 2a). Experimentally, high diffusivity 2*D* ∼ 1 *µm*^2^/*s* has been reported for higher buffer ionic strengths (19, 41) (Fig. 2b), and typically associated with a lack of apparent cohesin extrusion activity (37). The JDM therefore explains the apparent inhibition of cohesin extrusion on DNA *in vitro* in high-salt assays in terms of accelerated slippage kinetics.

**Figure 2.**
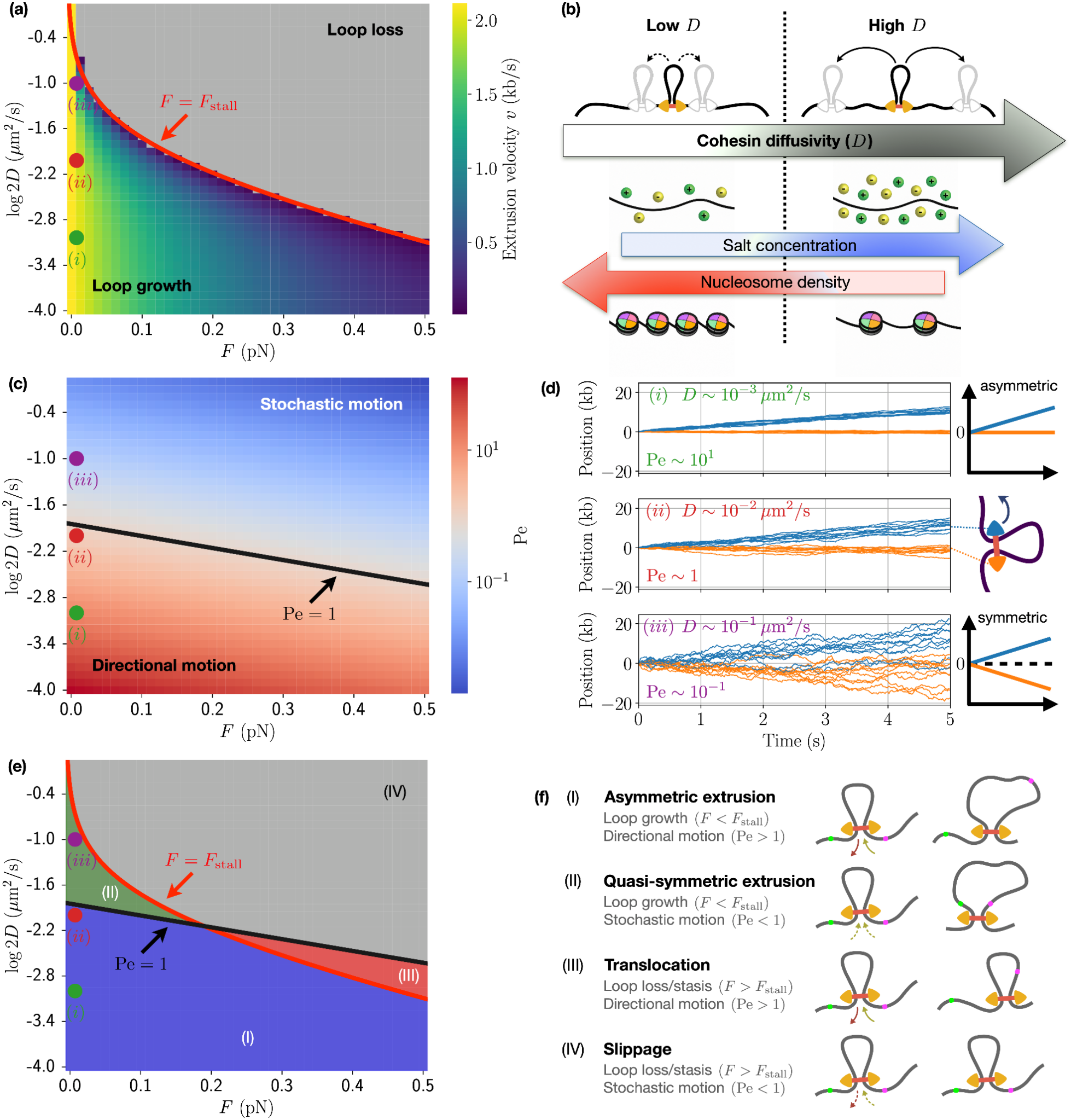
Cohesin behavior is determined by buffer and substrate conditions. **(a)** Cohesin extrusion velocity (*v*, Eq. (5)) as a function of applied DNA tension (*F*) and diffusivity (*D*). The red line indicates the diffusivity dependence of the cohesin stall force *F*_*stall*_, which delineates the regimes of loop growth and loop loss. **(b)** Respective impacts of buffer and substrate conditions on cohesin diffusivity *D*. **(c)** Péclet number (*Pe*, Eq. (9)), quantifying the relative roles of advective and diffusive transport on cohesin motion. The transition from stochastic to directional behavior occurs around *Pe* ∼ 1 (black line). **(d)** Single-cohesin anchor kinetics for *F* → 0 at low *(i)*, baseline *(ii)* and high *D (iii)*, as highlighted in Figs. 2a-b. Thin lines denote individual traces computed from stochastic simulations. **(e)** Kinetic phase diagram of cohesin showing: (I) asymmetric extrusion, (II) quasi-symmetric extrusion, (III) translocation, and (IV) slippage, where phase boundaries are shown as lines defined by the stall force *F*_*stall*,_ and the Péclet threshold. **(f)** Illustrations of kinetic behaviors I-IV, shown before and after a time update. Green and purple dots indicate reference positions along DNA as guides to the eye.

Additionally, the JDM predicts that low diffusivity substantially increases both cohesin stall force *F*_*stall*_ and mean extrusion velocity at fixed tension *F* < *F*_*stall*_ (Fig. 2a). Such lower diffusivity could result from lower ionic strengths, consistent with the higher stall forces and faster kinetics of cohesin extrusion reported in lower-salt buffer (10). Reduced diffusivity has also been reported upon the inclusion of nanometer-sized proteins and nucleosomes onto the DNA substrate (19) (Fig. 2b). Intriguingly, the JDM thus reveals that physical obstacles that reduce cohesin diffusion could in fact facilitate loop extrusion (Fig. 2a; see Discussion). Mechanistically, this occurs by inhibiting loop slipping while not significantly impeding active DNA capture (Eq. (5)). This possibility aligns with the reported ability of cohesin (10) and other SMCs (48–50) to bypass isolated nucleosomes, and indicates that nucleosomes at physiological densities could actually promote extrusion kinetics *in nucleo* (see Discussion).

Due to its differential impact on slippage versus capture, diffusivity determines the apparent directionality of cohesin motion. We quantified this via the Péclet number (51)

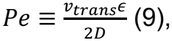

which interpolates between purely stochastic (*D* → ∞, *Pe* → 0) and purely directional behavior (*D* → 0, *Pe* → ∞), with a Péclet threshold *Pe* ∼ 1 typically separating the two (Fig. 2c). The degree of directionality sets the *apparent symmetry* of cohesin extrusion, as can be quantified in microscopy assays.

Experimentally, asymmetric extrusion is characterized by incorporation of DNA only from one side at a time into the loop (Fig. 2d). Quantified analogously, the JDM predicts that higher diffusivity increases the apparent symmetry of extrusion, as the directionality from asymmetric stepping is gradually concealed by the stochastic diffusion of the complex (Figs. 2c-d).

To quantify the impact of diffusivity on stochastic trajectories at the single-molecule level, we incorporated the JDM within a kinetic Monte-Carlo scheme (see Methods). We find that increased diffusivity gradually obscures the difference in kinetics effectively measured at the active and passive anchors in individual cohesin traces (Fig. 2d). This highlights the challenge for inferring intrinsic motor activity from single-molecule tracking data, and could account for the enhanced apparent symmetry of extrusion reported at higher salt for monomeric SMCs (16).

Using both the stall force and Péclet thresholds, apparent cohesin kinetics from single-molecule traces can be partitioned into four distinct regimes: (i) asymmetric extrusion, showing one-sided loop growth (low diffusion, low force); (ii) quasi-symmetric extrusion, showing apparent two-sided loop growth (high diffusion, low force); (iii) translocation, showing loop shrinkage or stasis with directional displacement (low diffusion, high force); and (iv) slippage, featuring loop shrinkage or stasis with stochastic motion (high diffusion, high force) (Figs. 2e-f). Remarkably, these four distinct regimes share a common coexistence point (*F* ≃ 0.2 *pN*, 2*D* ≃ 0.005 *µm*^2^/*s*), close to the forces and diffusivities reported for typical buffer conditions *in vitro* (*F* ∼ 0.1 *pN*, 2*D* ∼ 0.01 *µm*^2^/*s*). This close proximity to diverse kinetic regimes helps explain the range of extrusion behaviors reported across experimental assays. For instance, initial reports of symmetric extrusion by cohesin (10–13) could simply reflect small variations in DNA tension and cohesin diffusivity (i.e., buffer conditions) from more recent reports of asymmetric extrusion (14–17).

The JDM also provides insight into how physical aspects of assay setup can greatly impact observable kinetics. For example, experimental assays often use the force from buffer flow to stretch DNA in single-tethered curtain assays, but do not necessarily account for its impact during data analysis. In the JDM, buffer flow can be modeled as an approximately-linear tension gradient along the DNA contour (52). This position-dependent fiber tension in turn induces distinct behaviors with or against the flow (Fig. S2a). In the direction of flow, the JDM predicts that loops can grow at mean DNA tensions above the stall force measured in the absence of flow (Fig. S2b). This results from the preferential growth of cohesin loops towards the DNA free end, characterized by lower local tensions. The JDM can thus reconcile experimental observations of extrusion at tensions in flow-based assays (*F* > 0.5 *pN* (10)), well beyond reported cohesin stall forces at similar buffer compositions without flow (*F*_*stall*_ ∼ 0.15 *pN* (12, 14)).

Conversely, against the direction of flow the JDM predicts that the additional tension instead leads to purely translocative behavior (Fig. S2c), consistent with reports of cohesin translocation in single-tethered assays under strong flow conditions (11).

Together, these results demonstrate the ability of the JDM to quantitatively predict the sensitivity of cohesin extrusion to external physical and chemical cues, and provide mechanistic insights into the diversity of behaviors induced by subtle variations in experimental assay conditions.

### III. Cohesin mechanochemistry dictates its dynamic response to perturbations

Beyond its dependence on steady-state experimental conditions, cohesin activity also responds rapidly to mechanical and chemical perturbations. We thus applied the stochastic implementation of the JDM to characterize its predicted temporal responses to three perturbations: (i) changes in applied force, (ii) changes in buffer salt, and (iii) changes in cohesin ATPase activity.

We first considered JDM predictions upon abruptly increasing fiber tension beyond the cohesin stall force, as recently assayed experimentally using magnetic tweezers to apply a stretching force (34). We find that stretching-induced tension results in loop shrinkage preferentially from the passive anchor side (Fig. S3a), consistent with experimental observations (24). Asymmetry emerges because symmetric diffusive slippage is partially compensated by infrequent asymmetric capture steps at the active anchor, resulting in biased translocation kinetics towards the active anchor side. Importantly, asymmetric loop shrinkage does not by itself imply differential diffusivity of the two anchors, as it can arise solely from the underlying asymmetry of cohesin jumps (see Discussion).

We next analyzed JDM predictions after a sudden increase in cohesin diffusivity, mirroring the experimental effects of a high-salt wash (10, 11). Such experiments are typically performed in double-tethered assays, imposing a fixed end-to-end distance on the DNA fiber (Fig. S3b). In the JDM, a loop extruder on a fiber with a fixed end-to-end distance experiences self-limiting loop growth — DNA tension gradually increases up to an equilibrium loop size (Figs. 3a-b, S3b), which can be calculated analytically (see Methods). We then computed the response to a suddenly increased diffusivity from the baseline value 2*D* ≃ 0.01 *µm*^2^/*s* (*F*_*stall*_ ≃ 0.15 *pN*) (Fig. 3c). This analysis reveals a rapid loop shrinkage over the ∼10 s immediately following treatment and arrival at a new steady-state loop size with a correspondingly lower stall force, which matches analytical predictions in the new buffer conditions (Eq. (6); Fig. S3c).

**Figure 3.**
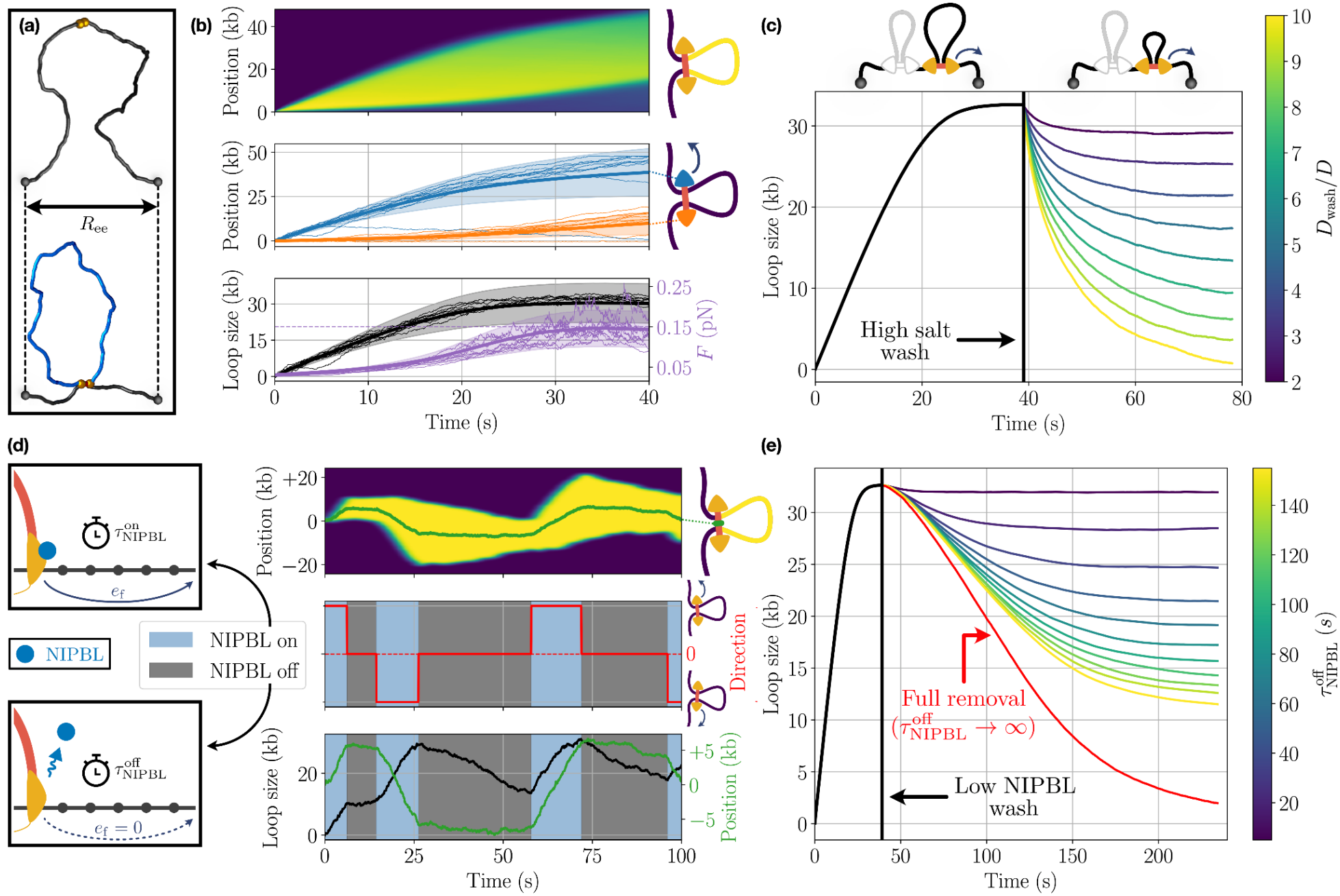
Cohesin mechanochemistry dictates its dynamic response to perturbations. **(a)** For double-tethered DNA assays, the JDM predicts that loop sizes grow until limited by increasing tension, and eventually reach a steady state upon approaching the stall force. Following panels show analyses for a double-tethered lambda-phage DNA substrate of length 48.5 *kb* with the two ends held at a fixed distance *R*_*ee*_ ≃ 3 *µm*. **(b)** *(top)* Average kymograph of loop growth, computed over 1000 independent stochastic traces loaded at the same initial reference position. *(middle)* Anchor positions over time (solid lines), displaying standard deviation (shading) and 10 illustrative traces (thin lines) with the active anchor in blue and passive in orange. *(bottom)* Same as above for the loop size (black) and DNA tension *F* (purple), showing characteristic steady-state plateau behavior for *F* → *F*_*stall*,_ = 0.15 *pN*. **(c)** Loop size versus time after suddenly increased diffusivity, mirroring the effect of a time-gated wash with a high salt buffer. Loop size rapidly decreases to a new steady-state value, which approaches 0 at very high salt. **(d)** Illustration of the stochastic binding kinetics of NIPBL as implemented within the JDM. When NIPBL is bound, extruders perform active steps. When NIPBL unbinds, extruder positions are driven solely by diffusion at both anchors. Representative traces over time show: (*top*) loop extent as a kymograph. (*middle*) direction (red) overlaid with periods of activity (blue) and inactivity (grey), displaying stochastic direction switching upon NIPBL rebinding (16). (*bottom*) loop size and position over time. (**e)** Loop size versus time after suddenly decreasing NIPBL concentration (modeled as the waiting time before NIPBL rebinding). Full removal (i.e., no rebinding) is shown in red.

These kinetics predicted by the JDM are consistent with the disappearance of loops observed upon high-salt buffer injection (10, 11). However, the JDM shows rapid loop loss can arise solely from increased loop slipping and does not require cohesin dislodging or inhibited ATPase activity, as previously conjectured (11) (see Discussion).

The third perturbation we considered was for cohesin ATPase activity via modulation of NIPBL kinetics or ATP concentration. We implemented the stochastic binding kinetics of NIPBL within the JDM as a two-state cohesin model featuring distinct NIPBL-bound 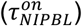 and NIPBL-free populations 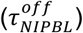. Residence time was set to match single-molecule measurements on loaded cohesin *in vitro* 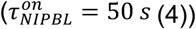. The cohesin jump rate was kept fixed for NIPBL-bound cohesins (*k* = 15.8 *s*^−1^, c.f. Sec. I), and was set to zero (*k* = 0) in the NIPBL-free state to reflect its lack of extrusion ability. To reproduce the stochastic switching of extrusion direction reported experimentally upon NIPBL reassociation (16), we further considered that cohesin passive and active anchors were randomly reassigned during each NIPBL binding event. Cohesins lacking NIPBL were described as two passive anchors, which both moved solely through diffusion. As such, the JDM predicts their motion displays tension-dependent loop loss (Fig. 3d) with a mean slippage velocity of ∼0.6 kb/s, closely matching experimental measurements of ∼0.4 kb/s in individual complexes lacking NIPBL on DNA *in vitro* (16) (c.f. Sec. I).

After instantaneous reductions in NIPBL concentration, final steady-state loop sizes were well predicted by an analytical mean-field expression for the effective stall force (Fig. S3d, see Supporting Information). Still, the loop shrinkage velocity upon NIPBL depletion was slower than upon high-salt wash because of its finite residence time on cohesin 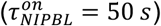, which effectively dampens the cohesin response upon rapid removal of free NIPBL from the buffer. Indeed, full NIPBL removal led to the disappearance of 95% of extruded loops over a timescale of ∼300 *s* (Fig. 3e), while less than 20 *s* were required to achieve 99% loop loss following high-salt injection (Fig. 3c). These slow model kinetics are consistent with experiments (97% loop dissolution after 480 *s* incubation in NIPBL-depleted buffer (11)), and differ from the predicted effects of ATP depletion — which was instead found to induce rapid (∼1min) loop shrinkage by directly inhibiting the active capture rate *k* (Figs. S3e-f). Thus, our analysis suggests that the dynamical association of cohesin accessory proteins also plays a crucial role in the regulation of loop extrusion mechanics.

Together, these analyses reveal how perturbations to mechanical conditions, buffer composition, or cohesin chemical state can all dynamically tune loop extrusion and generate highly-heterogeneous kinetics at the single-molecule level.

### IV. CTCF reduces local cohesin diffusion and inhibits active stepping via orientation-dependent pathways

Besides NIPBL association, cohesin extrusion is also tightly regulated by CTCF. *In vitro*, cohesin-CTCF encounters are highly transient and the majority display survival times below 3 minutes (14). Additionally, the ability of CTCF to act as a barrier to cohesin extrusion is not only polar — i.e., strongly conditioned on whether CTCF is encountered from its C- or N-terminal side — but also highly tension-dependent (14).

However, how both these experimental observations emerge mechanistically remains unknown. We thus implemented within the JDM a simple model of CTCF, in which we modeled DNA-bound CTCF as suppressing either cohesin diffusion and/or jump rates in a contiguous domain λ_*CTCF*_ ≃ 10 *nm* (or ∼30 *bp*), corresponding to the approximate identified length of the CTCF zinc finger binding region (53), imposing lowered diffusion (*D*_*CTCF*_) and/or jump rates (*k*_*CTCF*_) throughout the domain. Since the cohesin step size ϵ ≃ 40 *nm* significantly exceeds λ_*CTCF*_, this implies that cohesin can potentially step anywhere within the domain, or even bypass CTCF altogether as occasionally observed experimentally (14).

We first compared JDM analytical predictions with *in vitro* measurements for the survival time distribution of individual CTCF-cohesin encounters, starting from an arbitrary initial encounter position within the CTCF domain (see Supporting Information). By sweeping a range of suppression levels for both diffusion and capture, we found that best agreement can be achieved for *k*_*CTCF*_ = 0 and *D*_*CTCF*_ = 0.04 *D* (Figs. 4a and S4a). These inferences predict that CTCF fully inhibits stepping activity, and slows down cohesin diffusion ∼25 fold relative to DNA *in vitro*. Notably, with complete suppression of active stepping, the JDM predicts that survival decays asymptotically as τ^−1^, as opposed to the exponential behavior predicted by standard theories considering a constant cohesin-CTCF dissociation rate (14). This inverse-power-law decay at long times is a striking, non-trivial feature of the experimental data (Fig. 4a), and constitutes a clear dynamical signature of cohesin’s diffusion-dependent escape from CTCF.

**Figure 4.**
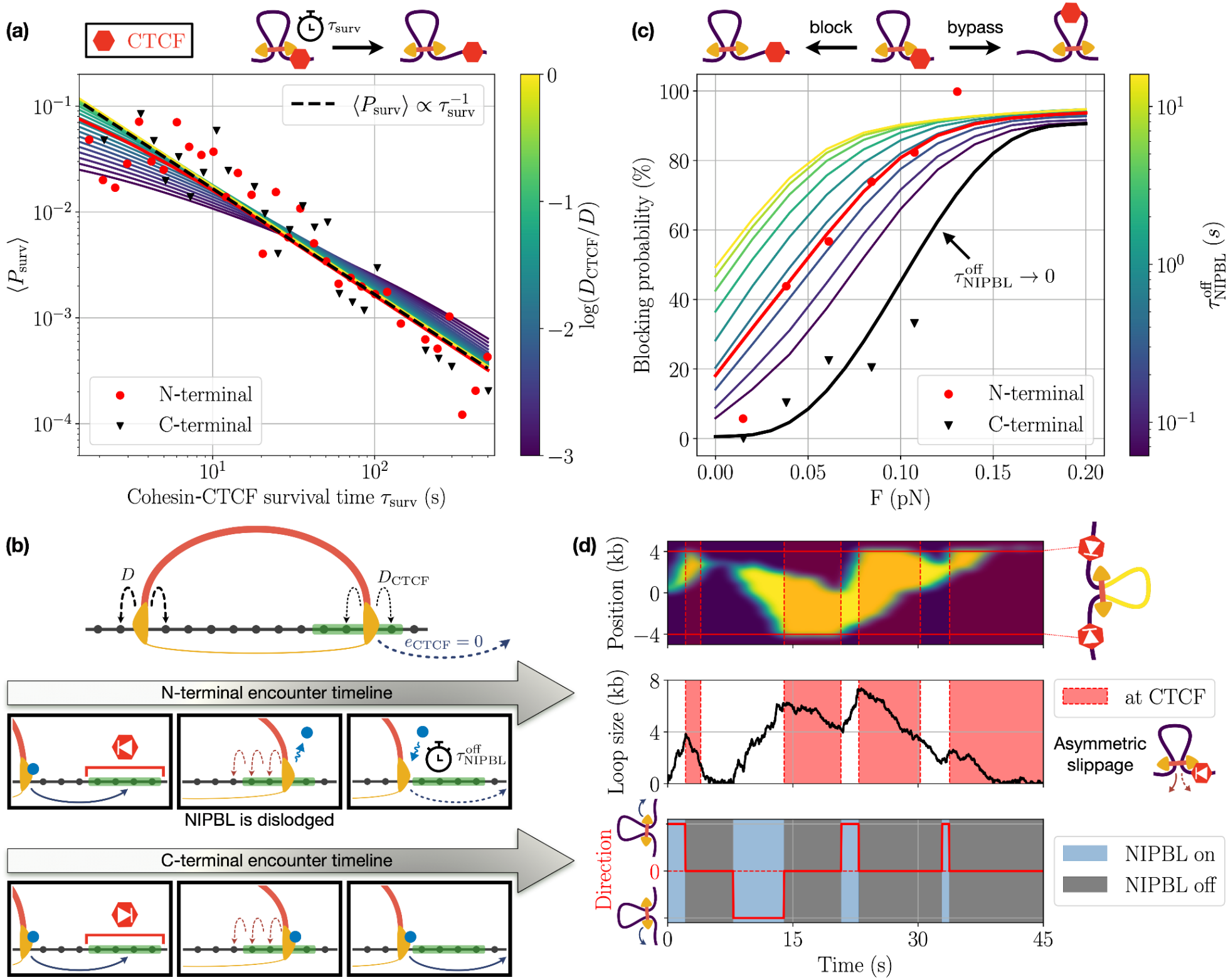
CTCF reduces local cohesin diffusion and inhibits active stepping via orientation-dependent pathways. **(a)** Survival time distribution of CTCF-cohesin associations, shown for experimental data (points, taken from (14)) and simulations (lines). Best agreement for both N- and C-terminal encounters (red line) shows fully inhibited active stepping and a ∼25 fold reduced diffusion (*D*_*CTCF*_ = 0.04 *D*) at CTCF (c.f. Fig. S4a). **(b)** Recapitulatory schematic of cohesin-CTCF encounter rules inferred by the JDM. While both C- and N-terminal encounters reduce local cohesin diffusivity and inhibit capture jumps, N-terminal encounters additionally impose a refractory period 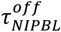 upon leaving CTCF before returning to an active NIPBL-associated state. **(c)** Blocking probability versus fiber tension for experimental data (points) and simulations (lines), as computed over a ∼1 minute simulation window. Best agreement is obtained for 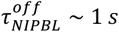 for N-terminal encounters (red line), and 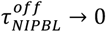 for C-terminal encounters (black). **(d)** Typical kymograph of N-terminal encounters, showing asymmetric loop slipping while cohesin is CTCF-bound, as well as stochastic switching in extrusion direction induced by NIPBL reassociation upon resolution of cohesin-CTCF contacts.

Importantly, agreement with survival kinetics alone does not distinguish whether CTCF suppresses stepping via direct inhibition of ATPase activity, modulation of cofactor association, or both. We thus investigated whether the JDM could discriminate between these mechanisms by comparison with additional experimental data. In this process we uncovered a puzzle: while the ability of CTCF to block extrusion seems highly polar *in vivo* (5, 6), experimental cohesin-CTCF survival kinetics are very similar for N- and C-terminal encounters *in vitro* (Fig. 4a) (14). The JDM therefore predicts that the local inhibition of cohesin diffusion and stepping by CTCF should occur in either encounter direction. We thus hypothesized that the difference in extrusion blocking ability exhibited by the CTCF N- and C-terminal could instead arise from differential orientation-dependent effects on cohesin chemical state. With this perspective informed by the JDM, *in vitro* observations where the cohesin direction switching frequency was reduced in buffer including CTCF N-terminal regions, but not C-terminal regions (17), could be consistent with a N-terminal-specific inhibition of NIPBL reassociation and/or promotion of its dissociation.

Thus, we considered a model in which N-terminal encounters imposed a refractory period 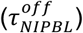 before stepping could resume following exit of cohesin from the CTCF domain. This refractory period could arise either from NIPBL dissociation from cohesin or its displacement from the active DNA strand by an N-terminally oriented CTCF encounter (Fig. 4b). We then assessed the ability of cohesin to bypass an isolated CTCF site as a function of DNA tension for a range of 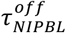. Comparison with available experimental data for N-terminal encounters yielded an optimal fit of 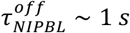, corresponding to rapid rebinding kinetics consistent with the high NIPBL stoichiometric excess of the extrusion assays (14) (Fig. 4c). Conversely, for C-terminal encounters, model predictions best aligned with experimental measurements in the limit 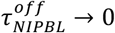, indicating the immediate availability of NIPBL on cohesin upon exit from the CTCF domain (Fig. 4c). These results provide insight into the origin of CTCF polarity: although C- and N-terminal encounters similarly inhibit cohesin diffusion and stepping kinetics, C-terminal encounters do not impose a refractory period for NIPBL-associated stepping. In contrast N-terminal encounters result in a transient refractory period before extrusion activity is reestablished (Fig. 4b; Figs. S4b-c).

The JDM additionally predicts that loops are not static over the course of CTCF-cohesin encounters *in vitro*, but rather display asymmetric slippage kinetics resulting in gradual loop loss at the distal cohesin anchor (Fig. 4d). This effect may be attributed to the faster diffusion kinetics — and hence, higher slippage — of the cohesin anchor away from CTCF, resulting in a tension-dependent loop shrinkage at typical velocity ∼0.5 *kb*/*s* while stepping is inhibited. This loop slipping behavior is predicted to be largely identical for N- and C-terminal encounters, given that the local inhibition of cohesin diffusion and stepping are independent of CTCF orientation in the JDM, and closely mirrors experimental measurements of loop shrinkage kinetics at CTCF *in vitro* (14). Furthermore, in the case of C-term-oriented bypass events, this diffusion asymmetry similarly leads to suppressed slippage at the passive anchor while the active anchor extrudes past CTCF, resulting in a higher net extrusion velocity *v* ≃ *v*_*active*,_ ∼ 1.3 *kb*/*s* as long as the passive anchor remains at CTCF (Eqs. (5) and (7); Fig. S4c). This value is consistent with the increased velocities *v* ∼ 1.42 *kb*/*s* reported experimentally after C-term encounters (54), thus providing a simple interpretation for the seemingly paradoxical ability of CTCF to transiently promote cohesin extrusion kinetics.

The role of diffusion for resolving cohesin encounters with CTCF led us to ask how CTCF blocking depends on cohesin diffusivity in the surrounding region. *In vivo*, CTCF binding sites are directly flanked by nucleosome-depleted regions (λ_*flank*_ ≃ 100 *bp*), which are in turn surrounded by arrays of well-positioned nucleosomes (55, 56). To investigate the functional roles of this effect using the JDM, we appended to each individual CTCF site two flanking regions of length λ_*flank*_ within which the local cohesin diffusion constant was set to *D*_*flank*_, and systematically varied *D*_*flank*_ from our baseline value *D* ≃ 0.005 *µm*^2^/*s* for DNA *in vitro* in extrusion buffer (c.f. Sec. I) to 0.01 *D* (Fig. S4d). This latter value corresponds to the estimated order of magnitude on nucleosomal DNA in physiological conditions, assuming a typical repeat length of ∼200 bp (19) (see Discussion), thus providing an estimate for cohesin diffusion on chromatin far from CTCF. We found that increased diffusion induced by nucleosome depletion upstream and downstream of CTCF led to a massive increase in CTCF blocking ability, with the local fiber tension required to reach 80% cohesin blocking during N-terminal encounters dropping from *F* ≃ 0.7 *pN* for *D*_*flank*_/*D* = 0.01 to *F* ≃ 0.3 *pN* for *D*_*flank*_/*D* = 1. This bolstered blocking simply emerges from increased slippage — and thus, reduced extrusion kinetics — in the vicinity of CTCF, and provides a mechanistic basis for the role of local chromatin remodeling in the regulation of cohesin-CTCF interactions.

Collectively, JDM analysis indicates that CTCF inhibits passive cohesin diffusion and suppresses active stepping via direction-dependent and independent pathways, resulting in blocking behavior that can be further modulated by the flanking chromatin context.

## Discussion

In summary, we presented a minimal biophysical description of cohesin that can be analytically parametrized using experimentally-measured mesoscopic quantities. This jump-diffusion model (JDM) represents cohesin motion as a competition between active DNA capture — characterized by discrete, directional jumps — and passive DNA slippage, which arises from the stochastic diffusion of the two cohesin anchors. This interplay is highly sensitive to applied tension, and gives rise to extrusion, translocation and loop slipping phases, thus rationalizing the full breadth of kinetic behaviors for cohesin reported *in vitro*. Such kinetic behaviors collectively emerge from the balance of active growth and passive shrinkage, revealing that the establishment and maintenance of cohesin loops are both dynamically regulated.

The JDM is based on a simple depiction of cohesin as an asymmetric ‘stepper’ featuring a pair of active and passive anchors, and is agnostic to the detailed molecular mechanisms underlying DNA capture and slippage. Our framework thus directly relates emergent cohesin dynamics at the mesoscale with their intrinsic motor kinetics at the single-molecule level. This provides understanding of SMC dynamics without requiring explicit modeling of the exact loading and active stepping process, as required by more detailed descriptions like the “swing-and-clamp” (33) or “hold-and-feed” model (57). In the contexts of these models, the two anchors of the JDM represent the effective seats of interaction between DNA and the core SMC complex, which may involve multiple distinct molecular interfaces (33, 58).

A key result of the JDM is the strong dependence of extrusion kinetics on cohesin diffusivity, which modulates both the rates of DNA slippage and the stochastic motion of the whole complex. Indeed, the low measured value of the stall force *in vitro* implies that observed loop loss kinetics are dominated by diffusion and that, at the range of experimentally probed forces, cohesin acts as a unidirectional motor. The rapid loop slipping induced by high diffusivity thus yields an overlooked explanation for the lack of evident cohesin extrusion reported in high-salt assays (19, 37, 41). Given its strong increase at higher salt, we envision that diffusivity reflects the affinity between DNA and the two cohesin anchors driven by electrostatics. While cohesin anchors could feature different diffusivities that bolster the asymmetry of cohesin kinetics, such differential diffusivities are not necessary to reproduce the asymmetric loop shrinkage reported upon DNA tension increase (24), which the JDM shows can be accounted for purely by residual, one-sided DNA capture (Fig. S3a). Furthermore, it suggests that rapid, salt-dependent loop loss does not necessarily argue against nontopological loading (11), but can emerge simply via increased slippage when weaker DNA-cohesin interactions bolster diffusivity. More generally, the JDM provides a simple interpretation for the strong susceptibility of cohesin loop maintenance to changes in experimental conditions, such as high-salt wash, NIPBL depletion or ATPase inhibition (10, 11), in terms of kinetic perturbations to cohesin slippage or capture that do not necessarily imply unloading the complex.

Even with a strictly one-sided intrinsic motor activity, the JDM predicts that the interplay between active stepping and stochastic diffusion can result in apparent extrusion kinetics that are either (quasi)symmetric or asymmetric. Importantly, transient changes in cohesin extrusion direction, as reported following NIPBL exchange (16) occur too infrequently to contribute to the apparent symmetry of a monomeric extruder over the experimentally-relevant timescales of minutes. Instead, we find that standard assay conditions for cohesin extrusion are near the boundary of quasi-symmetric versus asymmetric kinetic regimes. Thus, the JDM explains how seemingly conflicting reports of one-(14–17) and two-sided (10–13) extrusion by cohesin on naked DNA *in vitro* could be simply attributed to subtle changes in DNA tension and buffer conditions.

The quantitative relationship between stall force, substrate and buffer conditions predicted by the JDM has multiple important implications *in vivo*. First, loop diffusion (and consequently, slippage) should both be much slower on chromatin *in vivo* than observed on DNA *in vitro*, due to nucleosome inclusion on the substrate. Experimental observations indicate that the presence of nucleosomes at physiological densities would greatly suppress diffusion (∼1000-fold (19)), which would more than compensate for the higher slippage observed at physiological salt concentrations (∼150 mM vs ∼50 mM for extrusion assays on DNA *in vitro*). The conjunction of these effects is expected to reduce cohesin diffusivity ∼100-fold on native chromatin relative to typical extrusion assays (Dataset S1), which could be further regulated via differential nucleosome spacing depending on cellular and genomic context (59, 60).

Counterintuitively, the JDM predicts that reduced diffusivity on chromatin could promote extrusion due to reduced loop slipping. This provides a complementary pathway to recent proposals of boosted extrusion based on explicit models of nucleosome spacing (61, 62). Such detailed descriptions of nucleosome positions could be readily incorporated in the JDM to explore the local impacts of slippage versus occluded stepping on chromatin-based loop extrusion. The JDM additionally predicts that hindered passive motion would increase cohesin stall forces to ∼1 pN on chromatin, closer to that of well-characterized motor proteins such as myosin or kinesin (45, 46), as well as other physiological forces acting on chromatin such as protein-DNA co-condensation (∼0.2-0.8 pN (63)). Thus, the JDM reconciles the low cohesin stalling tensions reported on DNA *in vitro* with its ability to drive large-scale genome folding — and even potentially generate forces strong enough to shape overall nuclear morphology (64) — when loaded onto chromatin *in vivo*.

Besides the chromatin substrate, the JDM also predicts that cohesin loop extrusion kinetics *in vivo* are finely modulated by changes in nucleoplasmic composition, which may vary through the cell cycle or across cell types. For instance, increased slippage could explain how high-salt-based osmotic stress induces inhibition of topologically-associated domains and decrease of cohesin ChIP-seq signal in T47D (65), HeLa (66) and HCT116 cells (67), despite leaving loaded cohesin levels largely unaffected (65).

Such salt-induced slippage could also potentially bolster the loss of cohesin loops in early prometaphase, concomitant with a transient rise in intracellular Mg^2+^ ions (68). This could additionally facilitate their replacement by condensin loops (69), which the JDM would predict to be more salt resistant based on the slower basal diffusion kinetics of condensin (38).

The JDM enables insight into cohesin-CTCF encounters that are currently too challenging to analyze with more molecularly-detailed simulations. First, we find that CTCF locally inhibits both active extrusion and diffusion in either orientation. This yields a straightforward theoretical basis for why CTCF acts as a tension-mediated barrier: when extrusion is inhibited, tension strongly accelerates cohesin slippage away from CTCF, thus lowering bypass probabilities. Additionally, it provides a simple rationale for the increased extrusion rates observed following CTCF bypass events (54), which conversely results from inhibited slippage as the trailing passive anchor reaches CTCF (Fig. S4c). Second, polarity *in vitro* emerges via a distinct, N-terminal-mediated refractory period before extrusion can resume even after a cohesin-CTCF encounter has ended. This could reflect transient dislodging of NIPBL either from cohesin or the active DNA strand and could be promoted *in vivo* by cofactors not included in the analyzed reconstitutions (4), which could also modulate outcomes of C-terminal encounters (70). Thus, insights from the JDM motivate new single-molecule experiments tracking both CTCF and NIPBL to directly test for NIPBL eviction over N-terminal encounters and probe the timescale over which CTCF inhibits NIPBL re-binding, which are to our knowledge currently lacking.

The minimal assumptions of the JDM make it readily applicable to both monomeric and dimeric SMCs beyond cohesin. Indeed, monomeric condensin also diffuses along DNA in an ATP-independent fashion (38), albeit at lower rates than cohesin (*D* ∼ 10^−3^ *µm*^2^/*s* for condensin at ∼100 *mM* ionic strength, vs. *D* ∼ 10^−2^10^−1^ *µm*^2^/*s* for cohesin in similar conditions (Dataset S1)). Thus, in the JDM framework, the higher stall force (∼0.4 *pN* (18)) and stronger apparent asymmetry (12) reported for condensin could both be simply explained by reduced diffusivity. However, unlike cohesin, it is worth noting that condensin displays both a moderate force dependence in the forward capture step size as well as a non-negligible backwards stepping probability (22), which could both be tractably incorporated within the JDM (Eq. (7)) to provide a quantitative account of condensin stepping statistics. In assays with low cohesin diffusivity, the JDM predicts that active backward steps could also occur for cohesin at higher (> 1 *pN*) tensions probed with magnetic tweezers, which could account for occasional observations of reverse cohesin steps (32, 34). Moreover, dimeric SMCs could be treated as pairs of conjoined monomers, where symmetric capture kinetics naturally emerge from the mutualization of their two passive anchors domains. This includes SMC5/6 and Wadjet (15), and could additionally contribute to the apparent symmetry of cohesin extrusion based on occasional reports of cohesin dimerization *in vitro* (12, 15). Thus, the JDM provides a broadly-applicable mesoscale theory for loop extrusion by SMCs.

The JDM identifies important overlooked caveats of current reconstituted assays and their analysis. First, the strong predicted force dependence of extrusion rates (Fig. 1e) indicates that the reported range of values (11, 13) likely reflects poorly-constrained variations in tension across different experimental trajectories. Force-dependent kinetic measurements — similar to those reported for condensin (18) — would thus be required for more precise estimates of intrinsic motor dynamics. Second, apparent extrusion kinetics are sensitive to the cohesin diffusion rate, which is also widely distributed across experiments (Dataset S1). Beyond substrate and buffer, SMC diffusivity can also be increased by the inclusion of fluorescent tags (15). Such fluorescent tags are routinely employed in SMC diffusion assays (19, 40, 41), but can be absent in extrusion assays that track DNA alone (10–14). Conversely, the JDM also predicts that the lower DNA surface charge induced by the use of intercalating dyes in DNA tracking assays would result in increased diffusion — and thus, enhanced loop slipping — consistent with recent reports of accelerated condensin extrusion on dye-free DNA (71). Thus, the JDM highlights the importance of quantifying diffusion and extrusion under consistent experimental conditions to mitigate the effects of imaging biases on emergent SMC kinetics.

To conclude, the JDM provides a quantitative, first-principle theory for how local mechanical forces and chemical cues jointly govern cohesin loop extrusion over multi-minute timescales. While fully inferred from single-molecule experiments on DNA *in vitro*, the model also yields relevant insights into multiple *in vivo* cohesin regulatory pathways, and reveals how subtle changes in nucleoplasmic and chromatin environment may tune cohesin kinetics at the locus level. The parsimonious formulation of the JDM can be readily generalized to other SMCs, and can be tractably integrated within both polymer simulations of genome folding as well as previous network-based models of cohesin biochemistry (21). Our mechanistic framework thus bridges *in vivo* and *in vitro* observations of cohesin kinetics, and paves the way towards a comprehensive biophysical and chemical account of loop extrusion by SMCs.

## Materials and methods

### Stochastic simulation protocol

For stochastic simulations of the JDM, the rate Eqs. (1)-(3) were solved on a discrete 1D lattice with resolution δ = 2.5 *nm*. A total DNA contour length *l*_2_ = 48.5 *kb* (∼15 *µm*, or ∼6,000 sites) was considered to represent the full lambda phage genome, as employed in most reconstituted extrusion assays. Equations were integrated via a standard kinetic Monte-Carlo (MC) procedure with fixed timestep δ*t* = 1.5 µ*s*. This value was chosen to be low enough to yield a maximum diffusivity *D*_*max*_ = δ^2^/(2δ*t*) ∼ 2 *µm*^2^/*s*, in excess of the largest diffusivities *D* ∼ 1 *µm*^2^/*s* considered here. Cohesins were represented as a pair of anchors, each occupying one lattice position. Cohesin jumps were modelled by instantaneously displacing active anchors forward (i.e., in the direction of loop growth) by a distance ϵ = 40 *nm* (or 16 lattice sites) with rate *e*_*f*_. Asymmetric extrusion kinetics were implemented by setting *e*_*f*_ = 0 at passive anchors. Forward and backward diffusion steps were attempted by iterating anchor positions by a single site δ in either direction with respective rates *d*_*f,b*_. Each MC step consisted of independent jump and diffusive trial moves for each of the two cohesin anchors. NIPBL kinetics were described via association/dissociation moves with respective rates 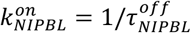 and 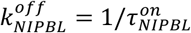 (Sec. III). In this case, NIPBL unbinding was modelled by treating both anchors as inactive, while NIPBL binding events resulted in stochastically switching one of the two anchors to the active state. Trial moves resulting in cohesin anchors stepping beyond these finite lattice boundaries or leading to any anchor overlap were both systematically rejected.

### Tension calculations in single- and double-tethered assays

For simulations of extrusion kinetics in single-tethered assays with external flow, we considered a linear tension gradient resulting in a contour-dependent force profile (Fig. S2a). Denoting by *s*_1_ and *s*_2_ the linear positions of the two cohesin anchors, we then applied the simulation protocol above using differential rates *e*_*f*_(*F*_1,2_) and *d*_*f,b*_(*F*_1,2_) at each anchor, with *F*_1,2_ = *F*(*s*_1,2_) the corresponding local DNA tension.

Conversely, for double-tethered assays, we instead assumed a fixed end-to-end distance *R*_*ee*_ < *l*_*c*_, and computed the length *l* of DNA outside the loop at each timestep via *l* = *l*_*c*_ – *s*_2_ + *s*_1_. We then obtained the mean, time-dependent DNA tension *F* as in experiments via the standard worm-like chain formula (10, 12, 14)

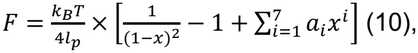

where *x* ≡ *R*_*ee*_/*l* is the relative DNA extension, *l*_*p*_ ∼ 50 *nm* the B-DNA persistence length and *a*_*i*_ a set of empirical numerical prefactors (72).

## Data and materials availability

Code describing simulations is publicly available: https://github.com/Fudenberg-Research-Group/discrete-time-extrusion/tree/jump_diffusion

## Acknowledgements

The authors thank Maxim Imakaev and Elphège Nora for helpful discussions and feedback, Xingchi Yan for feedback on the code, and all members of the Fudenberg group for discussions. MMCT was supported by the fellowship grant AWD-00010609 from the Gordon and Betty Moore Foundation and both authors were supported by NIGMS R35GM143116 to GF.

## Author Contributions

MMCT and GF conceived of the project. MMCT developed analytical theories, implemented simulations, and performed analyses. MMCT drafted the manuscript with GF.

## Competing Interest Statement

The authors declare no competing interests.

## Notes

### Competing Interest Statement

The authors have declared no competing interest.

## References

1. C. Hoencamp, B. D. Rowland, Genome control by SMC complexes. Nat. Rev. Mol. Cell Biol. 24, 633–650 (2023).

2. J. N. Wells, T. G. Gligoris, K. A. Nasmyth, J. A. Marsh, Evolution of condensin and cohesin complexes driven by replacement of Kite by Hawk proteins. Curr. Biol. CB 27, R17–R18 (2017).

3. R. Shah, et al., Cohesin cofactor dosage sets the rate of loop extrusion, rendering genome folding tunable yet vulnerable to genetic disruption. Mol. Cell 86, 1635–1652.e8 (2026).

4. G. Wutz, et al., PDS5 proteins control genome architecture by limiting the lifetime of cohesin-NIPBL complexes. Mol. Cell 86, 1614–1634.e10 (2026).

5. A. S. Hansen, CTCF as a boundary factor for cohesin-mediated loop extrusion: evidence for a multi-step mechanism. Nucleus 11, 132–148 (2020).

6. M. S. van Ruiten, B. D. Rowland, On the choreography of genome folding: A grand pas de deux of cohesin and CTCF. Curr. Opin. Cell Biol. 70, 84–90 (2021).

7. G. Fudenberg, et al., Formation of Chromosomal Domains by Loop Extrusion. Cell Rep. 15, 2038–2049 (2016).

8. A. L. Sanborn, et al., Chromatin extrusion explains key features of loop and domain formation in wild-type and engineered genomes. Proc. Natl. Acad. Sci. U. S. A. 112, E6456–6465 (2015).

9. G. Fudenberg, N. Abdennur, M. Imakaev, A. Goloborodko, L. A. Mirny, Emerging Evidence of Chromosome Folding by Loop Extrusion. Cold Spring Harb. Symp. Quant. Biol. 82, 45–55 (2017).

10. Y. Kim, Z. Shi, H. Zhang, I. J. Finkelstein, H. Yu, Human cohesin compacts DNA by loop extrusion.Science 366, 1345–1349 (2019).

11. I. F. Davidson, et al., DNA loop extrusion by human cohesin. Science 366, 1338–1345 (2019).

12. S. Golfier, T. Quail, H. Kimura, J. Brugués, Cohesin and condensin extrude DNA loops in a cell cycle-dependent manner. eLife 9, e53885 (2020).

13. T. L. Higashi, G. Pobegalov, M. Tang, M. I. Molodtsov, F. Uhlmann, A Brownian ratchet model for DNA loop extrusion by the cohesin complex. eLife 10, e67530 (2021).

14. I. F. Davidson, et al., CTCF is a DNA-tension-dependent barrier to cohesin-mediated loop extrusion. Nature 616, 822–827 (2023).

15. B. Pradhan, et al., Symmetry of loop extrusion by dimeric SMC complexes is DNA-tension-dependent. [Preprint] (2024).

16. R. Barth, et al., SMC motor proteins extrude DNA asymmetrically and can switch directions. Cell 188, 749–763.e21 (2025).

17. R. Barth, et al., Two CTCF motifs impede cohesin-mediated DNA loop extrusion. Mol. Cell 85, 4304–4317.e9 (2025).

18. M. Ganji, et al., Real-time imaging of DNA loop extrusion by condensin. Science 360, 102–105 (2018).

19. J. Stigler, G. Ö. Çamdere, D. E. Koshland, E. C. Greene, Single-Molecule Imaging Reveals a Collapsed Conformational State for DNA-Bound Cohesin. Cell Rep. 15, 988–998 (2016).

20. F. Corsi, E. Rusch, A. Goloborodko, Loop extrusion rules: the next generation. Curr. Opin. Genet. Dev. 81, 102061 (2023).

21. M. M. C. Tortora, G. Fudenberg, The physical chemistry of interphase loop extrusion. Cell Genomics 6, 101098 (2026).

22. J.-K. Ryu, et al., Condensin extrudes DNA loops in steps up to hundreds of base pairs that are generated by ATP binding events. Nucleic Acids Res. 50, 820–832 (2022).

23. A. Bonato, et al., Spontaneously directed loop extrusion in SMC complexes emerges from broken detailed balance and anisotropic DNA search. Nucleic Acids Res. 53, gkaf725 (2025).

24. G. Pobegalov, et al., Symmetry and force response of cohesin loop extrusion are determined by diffusion of its motor and anchor domains. [Preprint] (2026). Available at: https://www.biorxiv.org/content/10.64898/2026.06.08.730904v1[Accessed 16 September 2026].

25. A. J. Pinto, et al., SMC Motor Proteins Operate at the Near-Minimal Forces for DNA Loop Extrusion. [Preprint] (2026). Available at: https://www.biorxiv.org/content/10.64898/2026.03.05.709531v2[Accessed 27 September 2026].

26. J. F. Marko, P. De Los Rios, A. Barducci, S. Gruber, DNA-segment-capture model for loop extrusion by structural maintenance of chromosome (SMC) protein complexes. Nucleic Acids Res. 47, 6956–6972 (2019).

27. S. K. Nomidis, E. Carlon, S. Gruber, J. F. Marko, DNA tension-modulated translocation and loop extrusion by SMC complexes revealed by molecular dynamics simulations. Nucleic Acids Res. 50, 4974–4987 (2022).

28. K.-W. Moon, D.-G. Kim, J.-K. Ryu, Anisotropic scrunching of SMC with a baton-pass mechanism. Commun. Biol. 7, 881 (2024).

29. L. Dooms, S. Nomidis, E. Carlon, S. Gruber, J. F. Marko, Obstacle bypassing and intrinsically asymmetric loop extrusion in the segment-capture model of Structural Maintenance of Chromosomes complexes. bioRxiv 2026.07.07.736920 (2026). 10.64898/2026.07.07.736920.

30. R. Takaki, A. Dey, G. Shi, D. Thirumalai, Theory and simulations of condensin mediated loop extrusion in DNA. Nat. Commun. 12, 5865 (2021).

31. A. Dey, G. Shi, R. Takaki, D. Thirumalai, Loop Extrusion Reversal by Condensin Motor is Mediated by Catch Bonds. [Preprint] (2026). Available at: https://www.biorxiv.org/content/10.64898/2026.05.01.722258v1[Accessed 8 October 2026].

32. R. Janissen, R. Barth, I. F. Davidson, J.-M. Peters, C. Dekker, All eukaryotic SMC proteins induce a twist of -0.6 at each DNA loop extrusion step. Sci. Adv. 10, eadt1832 (2024).

33. B. W. Bauer, et al., Cohesin mediates DNA loop extrusion by a “swing and clamp” mechanism. Cell 184, 5448–5464.e22 (2021).

34. G. Pobegalov, L.-Y. Chu, J.-M. Peters, M. I. Molodtsov, Single cohesin molecules generate force by two distinct mechanisms. Nat. Commun. 14, 3946 (2023).

35. T. E. Melby, C. N. Ciampaglio, G. Briscoe, H. P. Erickson, The symmetrical structure of structural maintenance of chromosomes (SMC) and MukB proteins: long, antiparallel coiled coils, folded at a flexible hinge. J. Cell Biol. 142, 1595–1604 (1998).

36. C. H. Haering, J. Löwe, A. Hochwagen, K. Nasmyth, Molecular Architecture of SMC Proteins and the Yeast Cohesin Complex. Mol. Cell 9, 773–788 (2002).

37. I. F. Davidson, et al., Rapid movement and transcriptional re-localization of human cohesin on DNA. EMBO J. 35, 2671–2685 (2016).

38. T. Terakawa, et al., The condensin complex is a mechanochemical motor that translocates along DNA. Science 358, 672–676 (2017).

39. N.-L. Tanasie, P. Gutiérrez-Escribano, S. Jaklin, L. Aragon, J. Stigler, Stabilization of DNA fork junctions by Smc5/6 complexes revealed by single-molecule imaging. Cell Rep. 41, 111778 (2022).

40. M. Kanke, E. Tahara, P. J. Huisin’t Veld, T. Nishiyama, Cohesin acetylation and Wapl-Pds5 oppositely regulate translocation of cohesin along DNA. EMBO J. 35, 2686–2698 (2016).

41. S. Glaser, M. I. Molodtsov, J. F. X. Diffley, F. Uhlmann, Replisome passage through the cohesin ring. Cell 188, 5618–5631.e14 (2025).

42. R. Phillips, J. Kondev, J. Theriot, H. Garcia, Physical Biology of the Cell (Garland Science, 2012).

43. W. A. Woyczyński, Diffusion Processes, Jump Processes, and Stochastic Differential Equations (CRC Press, 2022).

44. M. E. Fisher, A. B. Kolomeisky, The force exerted by a molecular motor. Proc. Natl. Acad. Sci. U. S. A. 96, 6597–6602 (1999).

45. A. B. Kolomeisky, M. E. Fisher, A simple kinetic model describes the processivity of myosin-v. Biophys. J. 84, 1642–1650 (2003).

46. S. Liepelt, R. Lipowsky, Kinesin’s network of chemomechanical motor cycles. Phys. Rev. Lett. 98, 258102 (2007).

47. E. J. Banigan, L. A. Mirny, Loop extrusion: theory meets single-molecule experiments. Curr. Opin. Cell Biol. 64, 124–138 (2020).

48. M. Kong, et al., Human Condensin I and II Drive Extensive ATP-Dependent Compaction of Nucleosome-Bound DNA. Mol. Cell 79, 99–114.e9 (2020).

49. B. Pradhan, et al., SMC complexes can traverse physical roadblocks bigger than their ring size. Cell Rep. 41, 111491 (2022).

50. H. W. Liu, et al., The SMC Hinge is a Selective Gate for Obstacle Bypass. Nat. Commun. 16, 10457 (2025).

51. S. Redner, A Guide to First-Passage Processes (Cambridge University Press, 2001).

52. G. A. King, A. S. Biebricher, I. Heller, E. J. G. Peterman, G. J. L. Wuite, Quantifying Local Molecular Tension Using Intercalated DNA Fluorescence. Nano Lett. 18, 2274–2281 (2018).

53. J. Yang, et al., Structures of CTCF-DNA complexes including all 11 zinc fingers. Nucleic Acids Res. 51, 8447–8462 (2023).

54. H. Zhang, et al., CTCF and R-loops are boundaries of cohesin-mediated DNA looping. Mol. Cell 83, 2856–2871.e8 (2023).

55. Y. Fu, M. Sinha, C. L. Peterson, Z. Weng, The insulator binding protein CTCF positions 20 nucleosomes around its binding sites across the human genome. PLoS Genet. 4, e1000138 (2008).

56. C. T. Clarkson, et al., CTCF-dependent chromatin boundaries formed by asymmetric nucleosome arrays with decreased linker length. Nucleic Acids Res. 47, 11181–11196 (2019).

57. I. A. Shaltiel, et al., A hold-and-feed mechanism drives directional DNA loop extrusion by condensin. Science 376, 1087–1094 (2022).

58. R. van Wee, et al., NIPBL and STAG1 enable loop extrusion by providing differential DNA-cohesin affinity. Proc. Natl. Acad. Sci. U. S. A. 122, e2514190122 (2025).

59. A. Valouev, et al., Determinants of nucleosome organization in primary human cells. Nature 474, 516–520 (2011).

60. V. B. Teif, et al., Genome-wide nucleosome positioning during embryonic stem cell development. Nat. Struct. Mol. Biol. 19, 1185–1192 (2012).

61. A. Maji, R. Padinhateeri, M. K. Mitra, The Accidental Ally: Nucleosome Barriers Can Accelerate Cohesin-Mediated Loop Formation in Chromatin. Biophys. J. 119, 2316–2325 (2020).

62. N. Pellet, et al., Understanding the role of chromatin accessibility and compaction on loop extrusion dynamics. [Preprint] (2026). Available at: https://www.biorxiv.org/content/10.64898/2026.10.12.758711v1[Accessed 26 October 2026].

63. T. Quail, et al., Force generation by protein–DNA co-condensation. Nat. Phys. 17, 1007–1012 (2021).

64. I. Patta, et al., Nuclear morphology is shaped by loop-extrusion programs. Nature 627, 196–203 (2024).

65. R. Amat, et al., Rapid reversible changes in compartments and local chromatin organization revealed by hyperosmotic shock. Genome Res. 29, 18–28 (2019).

66. A. K. Golov, A. V. Golova, A. A. Gavrilov, S. V. Razin, Sensitivity of cohesin–chromatin association to high-salt treatment corroborates non-topological mode of loop extrusion. Epigenetics Chromatin 14, 36 (2021).

67. Y. Liu, J. Dekker, CTCF-CTCF loops and intra-TAD interactions show differential dependence on cohesin ring integrity. Nat. Cell Biol. 24, 1516–1527 (2022).

68. K. Maeshima, et al., A Transient Rise in Free Mg2+ Ions Released from ATP-Mg Hydrolysis Contributes to Mitotic Chromosome Condensation. Curr. Biol. CB 28, 444–451.e6 (2018).

69. K. Samejima, et al., Rules of engagement for condensins and cohesins guide mitotic chromosome formation. Science 388, eadq1709 (2025).

70. T. Földes, et al., Cohesin bridging as a physical principle of enhancer-promoter communication. [Preprint] (2026).

71. B. T. Analikwu, J. van der Torre, C. Dekker, Intercalator effects on DNA loop extrusion by SMC proteins. [Preprint] (2026). Available at: https://www.biorxiv.org/content/10.64898/2026.10.26.761753v1[Accessed 27 October 2026].

72. C. Bouchiat, et al., Estimating the persistence length of a worm-like chain molecule from force-extension measurements. Biophys. J. 76, 409–413 (1999).

