## Supplementary Information for "Interplay of diffusive and active motion governs cohesin extrusion and CTCF barrier function"

This PDF file includes:

- Supporting text
- Figs. S1 to S4
- Legends for Movies S1 to S3
- Legend for Dataset S1

Other supporting materials for this manuscript include the following:

- Movies S1 to S3
- Dataset S1

### Supporting Information Text

### Model derivation

Let us describe the cohesin loop extrusion process on a discrete, linear DNA lattice with arbitrarily-small genomic resolution $\delta$. We consider that the motion of cohesin arises from two distinct stochastic processes acting on each cohesin anchor: 1) microscopic (diffusion) steps of size $\delta$ to free adjacent lattice sites with rate $d$, and 2) mesoscopic (extrusion) jumps of size $\epsilon$ to distal target loci with rate $e$ (see Fig. 1a of the main text). The probability $P_i^{\mp}(t)$ to find a cohesin left ($-$) or right ($+$) anchor at site $i$ and time $t$ then satisfies the master equation

$$\frac{\partial P_i^{\mp}}{\partial t}(t) = P_{i-1}^{\mp}(t)d_{i-1,i}^{\mp} + P_{i+1}^{\mp}(t)d_{i+1,i}^{\mp} + P_{i-\epsilon}^{\mp}(t)e_{i-\epsilon,i}^{\mp} + P_{i+\epsilon}^{\mp}(t)e_{i+\epsilon,i}^{\mp} - P_i^{\mp}(t) \times \left[d_{i,i-1}^{\mp} + d_{i,i+1}^{\mp} + e_{i,i-\epsilon}^{\mp} + e_{i,i+\epsilon}^{\mp}\right]. \quad \text{[S1]}$$

Let us for now consider the case of an isolated cohesin motor, disregarding the presence of other potential local obstacles hindering its motion, and denote by $F$ the tension acting on each cohesin anchor. Although we here neglect the potential contour dependence of $F$ for the sake of simplicity, it is worth noting that $F$ is not necessarily uniform across the entire fiber, but should nonetheless display negligible variations across the cohesin jump size $\epsilon$ for the following discussions to remain valid. Assuming both anchors bear similar kinetics, we may write

$$\begin{aligned} d_{i,i-1}^{-} = d_{i+1,i}^{-} = d_{i,i+1}^{+} = d_{i-1,i}^{+} = d_{\mathrm{f}}(F), \\ d_{i-1,i}^{-} = d_{i,i+1}^{-} = d_{i+1,i}^{+} = d_{i,i-1}^{+} = d_{\mathrm{b}}(F), \\ e_{i,i-\epsilon}^{-} = e_{i+\epsilon,i}^{-} = e_{i,i+\epsilon}^{+} = e_{i-\epsilon,i}^{+} = e_{\mathrm{f}}(F), \\ e_{i-\epsilon,i}^{-} = e_{i,i+\epsilon}^{-} = e_{i+\epsilon,i}^{+} = e_{i,i-\epsilon}^{+} = e_{\mathrm{b}}(F), \end{aligned}$$

where $d_{\mathrm{f,b}}$ ($e_{\mathrm{f,b}}$) are the respective forward and backward diffusion (extrusion) step rates.

For clarity and without loss of generality, let us for now focus on the distribution of cohesin right anchors ($P_i^+$). The Taylor expansion of Eq. (S1) to second order in $\delta$ yields the standard Fokker-Planck equation for a stochastic jump-diffusion process,

$$\frac{\partial P_i^+}{\partial t} = -v_{\mathrm{l}}^+(F)\frac{\partial P_i^+}{\partial i} + D_{\mathrm{l}}^+(F)\frac{\partial^2 P_i^+}{\partial i^2} + e_{\mathrm{f}}(F) \times \left(P_{i-\epsilon}^+ - P_i^+\right) + e_{\mathrm{b}}(F) \times \left(P_{i+\epsilon}^+ - P_i^+\right), \quad \text{[S2]}$$

where the (local) drift velocity and diffusion coefficient $v_{\mathrm{l}}^+$ and $D_{\mathrm{l}}^+$ respectively read as

$$\begin{aligned} v_{\mathrm{l}}^+(F) &= \delta \times \left[d_{\mathrm{f}}(F) - d_{\mathrm{b}}(F)\right], \\ D_{\mathrm{l}}^+(F) &= \delta^2 \times \frac{d_{\mathrm{f}}(F) + d_{\mathrm{b}}(F)}{2}. \end{aligned} \quad \text{[S3]}$$

Note that in deriving Eq. (S2), we neglected the dependence of the cohesin extrusion jump size $\epsilon$ on the applied force $F$, consistent with the weak variations reported in single-molecule magnetic tweezer experiments, so that the force dependence of the extrusion jump kinetics may be fully captured by the coefficients $e_{\mathrm{f,b}}(F)$.

The mechanical work necessary to reel a DNA segment of length $\delta$ into the loop against the applied tension $F$ simply reads as $F\delta$. In the so-called Bell model of molecular motors [44], this work is split into two distinct kinetic contributions: 1) a fraction $\alpha F\delta$ which serves to inhibit forward steps (i.e. loop growth) via $d_{\mathrm{f}}$, and 2) a fraction $(1-\alpha)F\delta$ which conversely promotes backwards steps (i.e. loop shrinkage) via $d_{\mathrm{b}}$. Since $d_{\mathrm{f,b}}$ describe the passive, ATP-independent diffusion of individual cohesin anchors (i.e., $\Delta G = 0$), we may write

$$\begin{aligned} d_{\mathrm{f}}(F) &= k_{\mathrm{dif}} \times \exp\left(-\alpha\beta F\delta\right), \\ d_{\mathrm{b}}(F) &= k_{\mathrm{dif}} \times \exp\left[(1-\alpha)\beta F\delta\right], \end{aligned} \quad \text{[S4]}$$

where $\beta \equiv 1/k_{\mathrm{B}}T$. Eq. (S3)–Eq. (S4) thus yield, to second order in $\delta$,

$$v_{\mathrm{l}}^+(F) \equiv v_{\mathrm{slip}}^+(F) = -\delta^2 \times k_{\mathrm{dif}}\beta F + \mathcal{O}\left(\delta^3\right), \quad \text{[S5]}$$

$$D_{\mathrm{l}}^+(F) \equiv D = \delta^2 \times k_{\mathrm{dif}} + \mathcal{O}\left(\delta^3\right), \quad \text{[S6]}$$

irrespective of $\alpha$. Eq. (S5) and Eq. (S6) thus yield the fluctuation-dissipation relationship $v_{\mathrm{slip}}^+(F) = -\beta F D$ (Eq. (2) of the main text). Thus, Eq. (S5) and Eq. (S6) describe the ATP-independent component of cohesin motion, which is purely diffusive at 0 force ($v_{\mathrm{slip}}^+(F=0) = 0$) and increasingly biased towards loop slippage under finite applied loads ($v_{\mathrm{slip}}^+(F>0) < 0$).

In the case of active, ATP-dependent jumps, let us use the so-called *diffusive approximation* of jump-diffusion processes to similarly perform the Taylor expansion of Eq. (S2) to order $\epsilon^2$,

$$\frac{\partial P_i^+}{\partial t} = -v^+(F)\frac{\partial P_i^+}{\partial i} + D^+(F)\frac{\partial^2 P_i^+}{\partial i^2} + \mathcal{O}\left(\epsilon^3\right), \quad \text{[S7]}$$

where

$$\begin{aligned} v^+(F) &= v_{\mathrm{slip}}^+(F) + \epsilon \times \left[e_{\mathrm{f}}(F) - e_{\mathrm{b}}(F)\right], \\ D^+(F) &= D + \epsilon^2 \times \frac{e_{\mathrm{f}}(F) + e_{\mathrm{b}}(F)}{2}. \end{aligned} \quad \text{[S8]}$$

Identical considerations for the left anchor distribution $P_i^-$ similarly yield $v^-(F) = -v^+(F)$, while the independence of cohesin diffusion kinetics on ATP [10, 37] impose that the active component of cohesin diffusibility in Eq. (S8) is negligible, i.e., $D^-(F) = D^+(F) \simeq D$.

By analogy with Eq. (S4), the forward and backward jump rates $e_{\mathrm{f,b}}(F)$ may be written in the general form

$$\begin{aligned} e_{\mathrm{f}}(F) &= k_{\mathrm{ext}}^{\mathrm{f}} \times \exp\big(-\alpha\beta F\epsilon\big), \\ e_{\mathrm{b}}(F) &= k_{\mathrm{ext}}^{\mathrm{b}} \times \exp\big[(1-\alpha)\beta F\epsilon\big], \end{aligned} \tag{S9}$$

in which $k_{\mathrm{ext}}^{\mathrm{f}}/k_{\mathrm{ext}}^{\mathrm{b}} = \exp(-\beta\Delta G_0)$ is now determined by the chemical energy $\Delta G_0$ required to perform a productive extrusion jump at 0 applied tension. For cohesin, using experimentally-measured values $F_{\mathrm{stall}} \simeq 0.15\,\mathrm{pN}$ and $\epsilon \simeq 40\,\mathrm{nm}$, the maximum mechanical work $F_{\mathrm{stall}}\epsilon \simeq 6\,\mathrm{pN\,nm}$ provided by a forward extrusion jump is significantly lower than the energy $|\Delta G_0| \simeq 50\,\mathrm{pN\,nm}$ released by an individual ATP binding/hydrolysis event. Thus, the discussions in Sec. I of the main text show that $e_{\mathrm{b}}/e_{\mathrm{f}} \ll 1$, implying that the backward jump probability is negligible. Eq. (S8) and Eq. (S9) then yield the velocity of active anchors in the form

$$v^+(F) = v_{\mathrm{slip}}^+(F) + v_{\mathrm{capture}}^+(F), \tag{S10}$$

with $v_{\mathrm{capture}}^+$ the active DNA capture velocity,

$$v_{\mathrm{capture}}^+(F) \equiv \epsilon \times e_{\mathrm{f}}(F) = \epsilon k \exp\big(-\alpha\beta F\epsilon\big), \tag{S11}$$

where we introduced the shorthand $k \equiv k_{\mathrm{ext}}^{\mathrm{f}}$ (Eq. (4) of the main text). Using Eq. (S5), Eq. (S10) and Eq. (S11), the net extrusion velocity of the full cohesin complex reads as

$$v(F) \equiv v^+(F) - v^-(F) = -2\beta F D + \epsilon k \exp\big(-\alpha\beta F\epsilon\big), \tag{S12}$$

where we exploited the asymmetry of cohesin extrusion kinetics to write $v^-(F) = v_{\mathrm{slip}}^-(F) = -v_{\mathrm{slip}}^+(F) = \beta F D$. Using $v(F_{\mathrm{stall}}) = 0$, Eq. (S12) directly leads to

$$\epsilon k = 2\beta F_{\mathrm{stall}} D \times \exp\big(\alpha\beta F_{\mathrm{stall}}\epsilon\big), \tag{S13}$$

from which Eq. (6) of the main text immediately follows.

### Role of DNA tension distribution

To assess the impact of DNA tension in population-averaged measurements of extrusion kinetics, we exploited the experimental distribution $P(R_{\mathrm{ee}})$ of end-to-end distances reported in double-tethered extrusion assays (Fig. S1a, inset; Ref. [11]). The effective tension $F(l, R_{\mathrm{ee}})$ at arbitrary extruded loop length $l$ and fixed $R_{\mathrm{ee}}$ may be obtained using Eq. (10) of the main text,

$$F(l, R_{\mathrm{ee}}) = \frac{k_{\mathrm{B}}T}{4l_p} \times \left\{ \frac{1}{[1 - x(l, R_{\mathrm{ee}})]^2} - 1 + \sum_{i=1}^{7} a_i x(l, R_{\mathrm{ee}})^i \right\}, \tag{S14}$$

where $x(l, R_{\mathrm{ee}}) \equiv R_{\mathrm{ee}}/(l_c - l)$ is the relative DNA extension, with $l_p \simeq 50\,\mathrm{nm}$ the DNA persistence length and $l_c \simeq 15\,\mu\mathrm{m}$ the contour length of the lambda-phage genome. Let us disregard the role of cohesin unloading, which occurs over typical timescales of 5-10 minutes significantly beyond those of standard tension-based experiments in reconstituted assays [16]. The conditional probability of observing loops of a size $l$ for a given DNA extension $R_{\mathrm{ee}}$, $P(l \mid R_{\mathrm{ee}})$ is then inversely proportional to their growth or shrinkage velocity, $|v[F(l, R_{\mathrm{ee}})]|$, i.e.,

$$P(l \mid R_{\mathrm{ee}}) \propto \frac{1}{|v[F(l, R_{\mathrm{ee}})]|}, \tag{S15}$$

with $v$ the extrusion velocity (Eq. (S12)). Eq. (S15) signifies that loops in the rapid growth regime (i.e., at high $v$/low $F$) are highly transient, and thus less likely to be observed at any given time.

The empirical tension distribution $P(F)$ may be estimated from Eq. (S14) by drawing end-to-end distances $R_{\mathrm{ee}}$ from the prior distribution $P(R_{\mathrm{ee}})$, and subsequently sampling loop sizes $l$ based on the posterior probability $P(l \mid R_{\mathrm{ee}})$ (Eq. (S15)). The resulting tension distribution $P(F)$ is reproduced in Fig. S1a, and displays only moderate deviations from uniform behavior in the moderate tension range $F = 0.02$–$0.13\,\mathrm{pN}$. Discrepancies in the low and high tension ranges reflect the bias towards larger, slowly-growing loops in Eq. (S15). Such extremal regimes respectively involve small loop sizes and vanishing extrusion rates, and are both likely challenging to quantify reliably due to experimental and technical noise. To assess their impacts, we evaluated the mean extrusion rate $\langle v \rangle_P$ obtained using the empirical distribution $P(F)$, compared against the reference value $\overline{v} = 1\,\mathrm{kb/s}$ obtained assuming a uniform tension distribution (Eq. (7) of the main text). Numerically integrating Eq. (S12), Eq. (S14) and Eq. (S15) via standard Monte-Carlo sampling yields $\langle v \rangle_P \simeq 0.95\,\mathrm{kb/s}$. This close agreement with $\overline{v}$ demonstrates the validity of the uniformly-distributed tension approximation, and hence the robustness of the parameter inference procedure outlined in Sec. I of the main text.

### Roles of NIPBL kinetics and ATP depletion

The ability of cohesin to perform active extrusion jumps is contingent on the presence of its co-factor NIPBL. Since NIPBL associates with cohesin only transiently, this implies that cohesin DNA capture and NIPBL binding kinetics should be intimately coupled. Let us denote by $f_{\mathrm{NIPBL}} < 1$ the fraction of loaded cohesins bound by NIPBL, which depends on the respective average times $\tau_{\mathrm{NIPBL}}^{\mathrm{on,off}}$ an individual complex spends in the NIPBL-associated or dissociated state,

$$f_{\mathrm{NIPBL}} = \frac{\tau_{\mathrm{NIPBL}}^{\mathrm{on}}}{\tau_{\mathrm{NIPBL}}^{\mathrm{on}} + \tau_{\mathrm{NIPBL}}^{\mathrm{off}}}. \tag{S16}$$

In a mean-field approximation, the average capture rate $\overline{k}_{\Delta\mathrm{NIPBL}}$ is limited by the reduced portion of time spent in the NIPBL-bound state, and simply reads as

$$\overline{k}_{\Delta\mathrm{NIPBL}} = k \times f_{\mathrm{NIPBL}}, \tag{S17}$$

with $k$ the capture rate of the NIPBL-bound state at 0 tension (Eq. (S11)). Plugging Eq. (S16) and Eq. (S17) into Eq. (S13) directly leads to an expression for the mean-field stall force at finite NIPBL availability (Fig. S3d; see Eq. (6) of the main text),

$$\overline{F}_{\mathrm{stall}}^{\Delta\mathrm{NIPBL}} = \frac{1}{\alpha\beta\epsilon} W_0\left(\frac{\alpha\epsilon^2 k}{2D} \times \frac{\tau_{\mathrm{NIPBL}}^{\mathrm{on}}}{\tau_{\mathrm{NIPBL}}^{\mathrm{on}} + \tau_{\mathrm{NIPBL}}^{\mathrm{off}}}\right), \tag{S18}$$

with $W_0$ the principal branch of the Lambert $W$ function.

Conversely, ATP depletion serves to directly inhibit the active jumping rate in the NIPBL-bound state $k_{\Delta\mathrm{ATP}} = k \times f_{\mathrm{ATP}}$, as described by a scaling factor $f_{\mathrm{ATP}}$ determined by the stoichiometry of ATP relative to the cohesin-NIPBL complex. The resulting reduced stall force $F_{\mathrm{stall}}^{\Delta\mathrm{ATP}}$ may thus be obtained by simply substituting $f_{\mathrm{ATP}}$ for $f_{\mathrm{NIPBL}}$ in Eq. (S18) (Fig. S3f),

$$F_{\mathrm{stall}}^{\Delta\mathrm{ATP}} = \frac{1}{\alpha\beta\epsilon} W_0\left(\frac{\alpha\epsilon^2 k}{2D} \times f_{\mathrm{ATP}}\right). \tag{S19}$$

### Role of CTCF-cohesin interactions

Let us now consider the presence of CTCF, which may affect the frequency of both cohesin diffusion steps and extrusion jumps in the JDM via distinct, position-specific prefactors $d^{i,\mathrm{CTCF}}$ and $e^{i,\mathrm{CTCF}}$. For simplicity, let us assume that these effects act uniformly within a genomic neighborhood of typical length $\lambda_{\mathrm{CTCF}}$ around a CTCF motif, such that for any site $i$ within the CTCF domain,

$$\begin{aligned} d_{\mathrm{f,b}}^{i,\mathrm{CTCF}}(F) &= d_{\mathrm{CTCF}} \times d_{\mathrm{f,b}}(F), \\ e_{\mathrm{f,b}}^{i,\mathrm{CTCF}}(F) &= e_{\mathrm{CTCF}} \times e_{\mathrm{f,b}}(F), \end{aligned} \tag{S20}$$

and such that $d_{\mathrm{f,b}}^{i,\mathrm{CTCF}} = d_{\mathrm{f,b}}$ and $e_{\mathrm{f,b}}^{i,\mathrm{CTCF}} = e_{\mathrm{f,b}}$ at loci $i$ outside the CTCF domain, to recover unimpeded cohesin kinetics on naked DNA. In Eq. (S20), $d_{\mathrm{CTCF}} \le 1$ and $e_{\mathrm{CTCF}} \le 1$ then correspond to damping factors describing hindered cohesin motion in the vicinity of CTCF, relative to the baseline rates $d_{\mathrm{f,b}}$ and $e_{\mathrm{f,b}}$ far away from CTCF (see Eq. (S4) and Eq. (S9)).

Under these assumptions, Eq. (S1) may be recast in the more general form

$$\frac{\partial P_i^{\mp}}{\partial t} = -\frac{\partial\left(v_i^{\mp} P_i^{\mp}\right)}{\partial i} + \frac{\partial^2\left(D_i^{\mp} P_i^{\mp}\right)}{\partial i^2} + \mathcal{O}\left(\epsilon^3\right), \tag{S21}$$

where the locus-dependent diffusion coefficient is such that $D_i^{\mp} = d_{\mathrm{CTCF}} \times D \equiv D_{\mathrm{CTCF}}$ within a CTCF domain, and simply reduces to $D_i^{\mp} = D$ away from CTCF. The net velocity $v_i^{\mp} \equiv v_{\mathrm{CTCF}}^{\mp}$ of an active cohesin anchor at CTCF then reads as (see Eq. (S12))

$$v_{\mathrm{CTCF}}^{\mp}(F) = \mp e_{\mathrm{CTCF}} \times \epsilon k \exp\left(-\alpha\beta F\epsilon\right) \pm d_{\mathrm{CTCF}} \times \beta F D. \tag{S22}$$

For simplicity and without loss of generality, let us now derive the probability for a cohesin right-anchor locating at an arbitrary initial position $x_0 \in [0, \lambda_{\mathrm{CTCF}}]$ within a CTCF domain to dissociate from CTCF without bypass, i.e., by exiting through the leftmost CTCF boundary, which we here set to position 0. Let us define the Péclet number of the cohesin anchor at CTCF (see Eq. (9) of the main text),

$$\mathrm{Pe}(F, x_0) \equiv \frac{v_{\mathrm{CTCF}}^{+}(F) \times x_0}{2D_{\mathrm{CTCF}}}. \tag{S23}$$

This probability may be identified as the first-passage probability $P_{\mathrm{first}}$ for cohesin at position 0 via the drift-diffusion process described by Eq. (S21)– Eq. (S23), which may be derived in the form [51]

$$P_{\mathrm{first}}(\tau, F, x_0) = \frac{1}{2}\left\{1 - \mathrm{erf}\left(\frac{x_0}{\sqrt{4D_{\mathrm{CTCF}}\tau}} + \Delta(\tau, F, x_0)\right) + e^{-2\mathrm{Pe}(F, x_0)}\left[1 - \mathrm{erf}\left(\frac{x_0}{\sqrt{4D_{\mathrm{CTCF}}\tau}} - \Delta(\tau, F, x_0)\right)\right]\right\}, \tag{S24}$$

where $\Delta(\tau, F, x_0) \equiv \mathrm{Pe}(F, x_0)\sqrt{D_{\mathrm{CTCF}}\tau}/x_0$ and erf is the Gauss error function. The average survival probability $\langle P_{\mathrm{surv}}\rangle(\tau)$ may then be obtained by integrating Eq. (S24) over accessible tensions $F$ and cohesin positions $x_0$ within the CTCF domain,

$$\langle P_{\mathrm{surv}}\rangle(\tau) = 1 - \frac{1}{F_{\mathrm{stall}}\lambda_{\mathrm{CTCF}}} \times \int_0^{F_{\mathrm{stall}}} dF \int_0^{\lambda_{\mathrm{CTCF}}} dx_0\, P_{\mathrm{first}}(\tau, F, x_0). \tag{S25}$$

Let us assume a typical footprint $\lambda_{\text{CTCF}} \simeq 10\,\text{nm}$ for an individual DNA-bound CTCF protein, and use extrusion parameters inferred previously for assays with naked DNAaway from the CTCF region (see Sec. I of the main text). Sweeping over potential values of $d_{\text{CTCF}}$ and $e_{\text{CTCF}}$, numerical integration of Eq. (S25) by standard quadrature methods yields optimal agreement with experimental data for $d_{\text{CTCF}} \simeq 0.04$ and $e_{\text{CTCF}} = 0$ (Fig. S4a and Fig. 4a of the main text), indicating that CTCF fully inhibits extrusion activity and slows down cohesin diffusion kinetics $\sim 25$-fold. In this extrusion-free regime, one may show using the asymptotic properties of the error function that Eq. (S24) reduces to, in the limit of long residence times $\tau$,

$$\lim_{\tau\to\infty} P_{\text{first}}(\tau, F, x_0) = 1 - \sqrt{\frac{4D_{\text{CTCF}}}{\pi}} \times \frac{x_0}{v^+_{\text{CTCF}}(F)^2 \tau^{3/2}} \times \exp\left[-\frac{v^+_{\text{CTCF}}(F)^2 \tau}{4D_{\text{CTCF}}}\right],$$

so that Eq. (S25) yields

$$\begin{aligned}\lim_{\tau\to\infty} \langle P_{\text{surv}}\rangle(\tau) &= \frac{\lambda_{\text{CTCF}} \times \Gamma(-1/2)}{8\sqrt{\pi} v^+_{\text{CTCF}}(F_{\text{stall}})} \times \frac{1}{\tau} \\ &= \frac{\lambda_{\text{CTCF}}}{4\beta D_{\text{CTCF}} F_{\text{stall}}} \times \frac{1}{\tau},\end{aligned} \qquad \text{[S26]}$$

with $\Gamma$ the Euler gamma function. Hence, the survival probability surprisingly decays asymptotically as $\tau^{-1}$, as opposed to the exponential behavior predicted by standard theories considering a constant cohesin-CTCF dissociation rate [14]. Such an inverse-power-law decay at long times is a striking feature of the experimental data reproduced in Fig. 4a of the main text. These slower survival statistics therefore suggest a significantly-greater relative prevalence of long-lived cohesin-CTCF association events than predicted by naive exponential-based descriptions, and arise from the slow, biased diffusion of cohesin anchors within finite-sized CTCF domains.

Interestingly, despite this long-tailed survival behavior, the majority of CTCF-cohesin co-localization events are predicted to be highly transient. Indeed, although the $\tau^{-1}$ asymptote implies that the time integral of $\langle P_{\text{surv}}\rangle$ diverges — and hence, that the mean cohesin residence time is theoretically unbounded — the average survival time within an arbitrary finite, experimentally-accessible measurement window $\tau_{\text{meas}} \sim 10\,\text{min}$ reads as

$$\langle \tau_{\text{res}}\rangle \equiv \int_0^{\tau_{\text{meas}}} d\tau \, \langle P_{\text{surv}}\rangle(\tau) \sim 1.2\,\text{s}. \qquad \text{[S27]}$$

Thus, the model predicts that cohesin-CTCF encounters are largely resolved over timescales of just a few seconds, consistent with single-molecule observations in reconstituted assays [14].

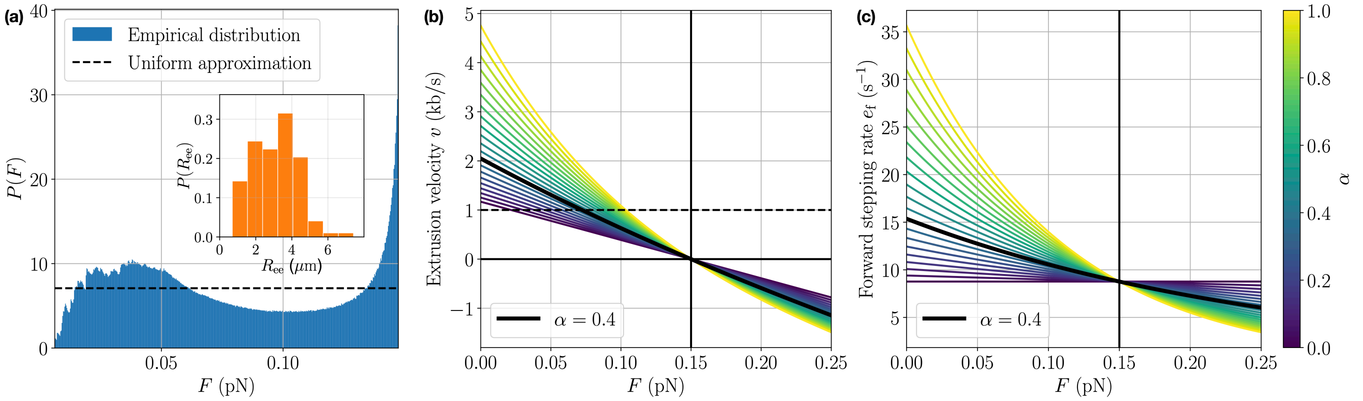


**Fig. S1. Jump-diffusion model parametrization & tension-dependent kinetics (related to Fig. 1).**
**(a)** Comparison of the empirical tension distribution, inferred from the experimental prior distribution of end-to-end tether distances ($R_{\mathrm{ee}}$, inset; from Ref. (11)), against the uniform approximation in Eq. (7) of the main text. **(b)-(c)** Tension dependence of the mean extrusion velocities (b) and cohesin stepping rates (c) in the JDM for different values of the load distribution parameter $\alpha$ (Eqs. (4)-(5)). The vertical line denotes the cohesin stall force $F_{\mathrm{stall}} = 0.15\,\mathrm{pN}$ on DNA in vitro, assumed as baseline in all curves. The dashed horizontal line indicates the corresponding mean extrusion velocity $\overline{v} = 1\,\mathrm{kb/s}$, averaged over accessible tensions $F < F_{\mathrm{stall}}$, which uniquely specifies the value $\alpha = 0.4$ for cohesin in the JDM.

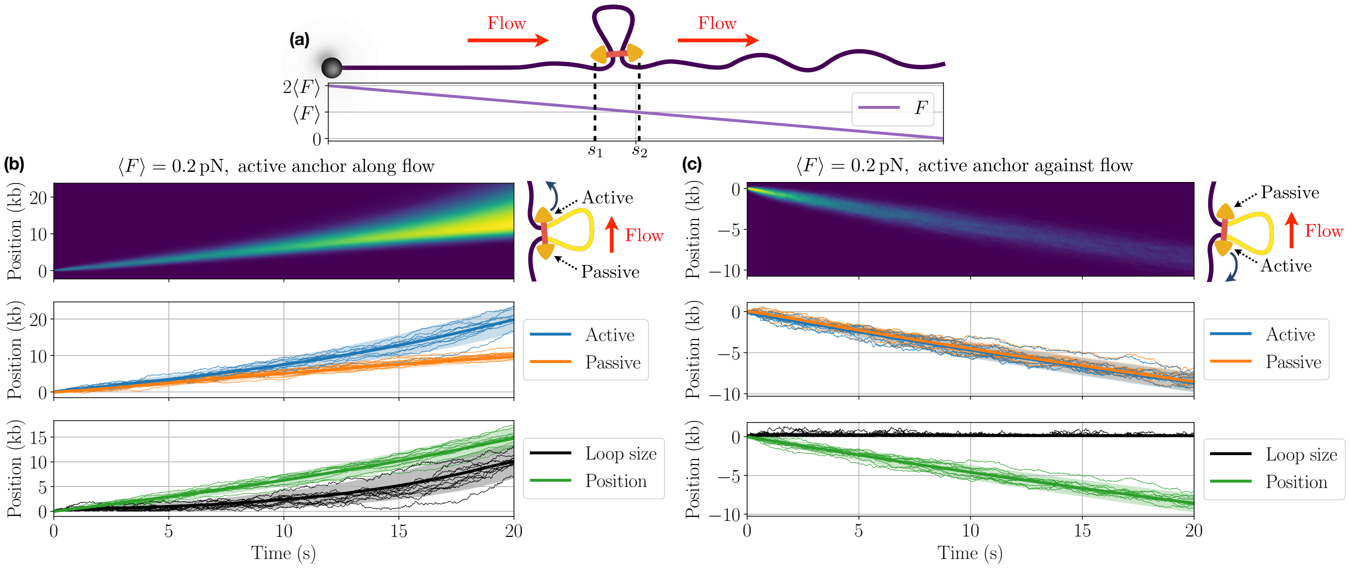


**Fig. S2. Active buffer flow in single-tethered extrusion assays affects cohesin kinetics (related to Fig. 2).**
**(a)** Sustained application of external flow, as employed in single-tethered DNA curtain experiments, results in an approximately-linear tension gradient directed towards the tethered DNA end. This effect can be captured in the JDM by substituting for the mean tension $F$ the local stretching forces $F_1 = F(s_1)$ and $F_2 = F(s_2)$ at the local positions $s_1 < s_2$ of the two cohesin anchors (Eq. (5) of the main text). **(b)** The lower tension near the free DNA end results in facilitated extrusion when active steps proceed along the flow direction (i.e., when $s_2$ marks the location of the active anchor). This results in the formation of cohesin loops even when the contour-averaged tension $\langle F\rangle = 0.2\,\mathrm{pN}$ exceeds the cohesin stall force ($F_{\mathrm{stall}} = 0.15\,\mathrm{pN}$). Thin lines denote individual traces, while solid lines and shading represent averages and standard deviations obtained over 1000 independent samples. **(c)** Conversely, extrusion is suppressed if stepping is directed against the flow, resulting in cohesin translocation without loop growth towards the tethered DNA end.

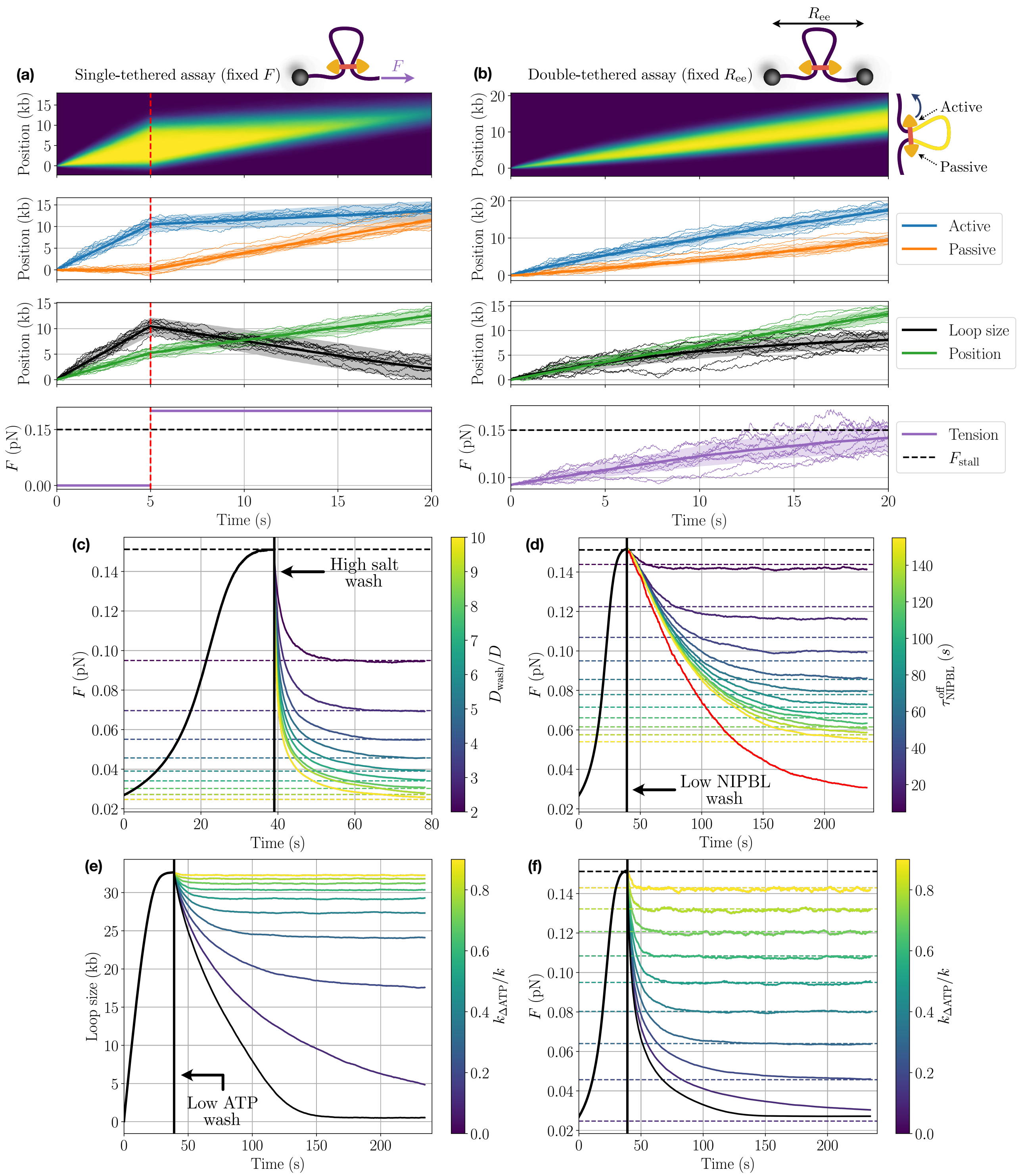


**Fig. S3. Loop kinetics are dynamically controlled by assay setup and applied forces (related to Fig. 3).**
**(a)** Abruptly increased tension (red dashed line at 5 seconds) in a singly-tethered assay results in loop loss preferentially from the passive anchor side. Panels from top to bottom: loop extent as a kymograph, motor and anchor positions, loop size and position, and applied force over time. Stochastic traces are obtained as in Fig. 2d. **(b)** A JDM extruder on DNA with a fixed end-to-end distance results in self-limiting loop growth up to a steady-state loop size. From top to bottom: loop extent as a kymograph, motor and anchor positions, loop size and position, and increasing force over time (dashed line shows the cohesin stall force, reached at final steady-state). **(c)** High salt wash in double-tethered cohesin assays results in rapid loop loss from increased slippage, until DNA tension reaches the new cohesin stall force $F_{\mathrm{stall}}^{\mathrm{wash}}$ (dashed lines) matching the higher cohesin diffusivity $D_{\mathrm{wash}}$ (Fig. 3b; see Eq. (6)). **(d)** Same as (c) following NIPBL depletion, showing slower loop shrinkage and relaxation towards a reduced stall force due to increasingly-long periods of cohesin inactivity, which is well-captured by mean-field theory (Eq. (S18)). Full removal (i.e., no rebinding) is shown in red. **(e)-(f)** ATP depletion conversely leads to a direct inhibition of the capture rate $k_{\Delta\mathrm{ATP}}$ in the active cohesin state, which results in rapid loop loss and decreased stall force (dashed lines, see Eq. (S19)).

 

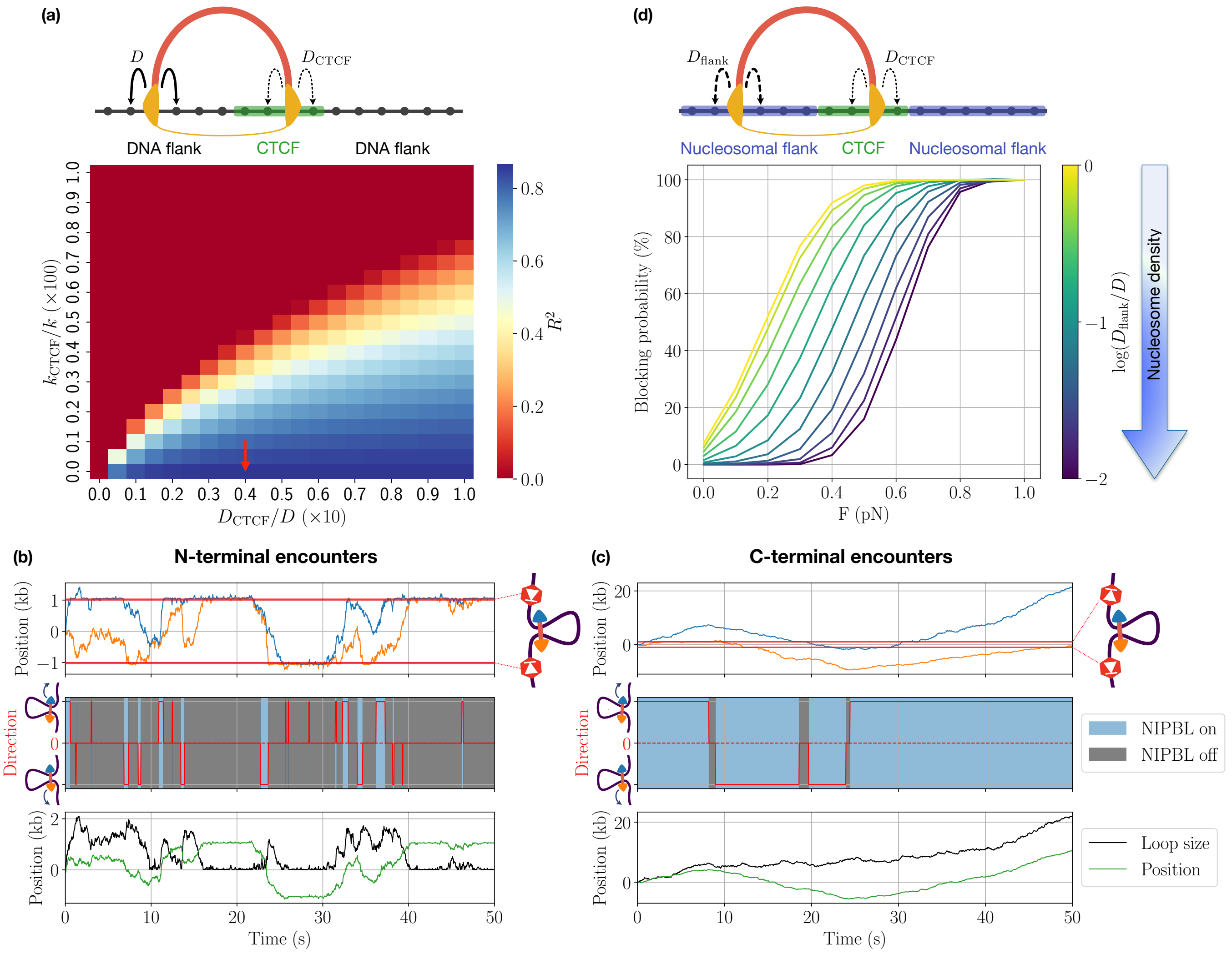


**Fig. S4. CTCF and flanking regions jointly affect cohesin kinetics (related to Fig. 4).**
**(a)** Systematic sweep of cohesin diffusion ($D_{\mathrm{CTCF}}$) and active step rate ($k_{\mathrm{CTCF}}$) at CTCF shows best agreement with experimental survival statistics measured in (14) for $k_{\mathrm{CTCF}} = 0$ and $D_{\mathrm{CTCF}} = 0.04\,D$ (red arrow; see Fig. 4a). **(b)** N-terminal oriented encounters lead to a refractory period, resulting in increased blocking efficiency and higher frequency of exchanges in the extrusion direction. **(c)** C-terminal encounters conversely have no effect on cohesin-NIPBL association, resulting in higher bypass probabilities and lower (baseline) direction switching. **(d)** Nucleosome inclusion in the CTCF flanking regions reduces blocking efficiency by locally inhibiting cohesin diffusivity and loop slipping, suggesting a potential regulatory role for chromatin remodeling at CTCF sites.

**Movie S1. Exemplary cohesin kinetic trace predicted by the JDM at low DNA tension ($F \simeq 0.05\,\mathrm{pN}$). Movie is slowed down x20 relative to real time. A linear DNA segment of length 1,600 base pairs is shown.**

**Movie S2. Same as Movie S1 at intermediary tension near the cohesin stall force ($F \simeq F_{\mathrm{stall}} = 0.15\,\mathrm{pN}$).**

**Movie S3. Same as Movie S1 at high tension ($F \simeq 0.25\,\mathrm{pN}$).**

**SI Dataset S1 (DatasetS1.xlsx)**

Curated list of cohesin diffusivities ($D$) and extrusion velocities ($v$) assayed across various reconstitution conditions *in vitro*, with corresponding literature references. "NR" indicates values not reported or quantified.
